# Discovery of novel compounds and target mechanisms using a high throughput, multiparametric phenotypic screen in a human neuronal model of Tuberous Sclerosis

**DOI:** 10.1101/2024.02.22.581652

**Authors:** Luis A. Williams, Steven J. Ryan, Vaibhav Joshi, Caitlin Lewarch, Amy Elder, Owen McManus, Patrice Godard, Srinidhi Sridhar, Jane Jacques, Jennifer Grooms, James J. Fink, Hongkang Zhang, Michel Gillard, Cécile Pegurier, Yogesh Sabnis, Véronique André, Lucinda Steward, Stefanie Dedeurwaerdere, Orrin Devinsky, Christian Wolff, Graham T. Dempsey

## Abstract

Tuberous sclerosis complex (TSC) is a rare genetic disorder caused by mutations in the mTOR pathway genes *TSC1* or *TSC2*. TSC can affect multiple organs including the brain, and most patients (75-90%) present with seizures during early childhood and intractable epilepsy throughout life. mTOR inhibitors, part of the current standard of care, lack the optimal characteristics to fully address patient phenotypes. Here, we report on the application of our all-optical electrophysiology platform for phenotypic screening in a human neuronal model of TSC. We used CRISPR/Cas9-isogenic *TSC2^−/−^* iPS cell lines to identify disease-associated changes to neuronal morphology, transcript expression and neuronal excitability. We established a robust multiparametric electrophysiological phenotype which we then validated in TSC patient-derived neurons. We used this phenotype to conduct a screen of ∼30,000 small molecule compounds in human iPS cell-derived neurons and identified chemical scaffolds that rescued the functional TSC disease parameters. Confirmed hits may act via different mechanisms than direct mTOR pathway inhibition. This strategy provides molecular starting points for therapeutic development in TSC and a framework for phenotype discovery and drug screening in other neurological disorders.

## INTRODUCTION

Successful drug development for central nervous system (CNS) disorders remains a challenge, with clinical failures outpacing successes^1^. Rare genetic diseases with identified genetic causes represent a tractable near-term opportunity for therapeutic development, with ∼70% of the ≥6,000 identified genetically defined disorders having an impact on the CNS^2,3^. Severe neurodevelopmental and seizure disorders represent a disease area of significant unmet medical need, often affecting patients early in life^2,3,4,5^. When considering non-lesion pediatric epilepsy patients, ∼25-45% have a known genetic cause with identified gene targets spanning diverse cellular functions including ion channels, synaptic proteins, mTOR (mammalian target of rapamycin) pathway regulators and chromatin remodeling and transcription regulators^3,4,5^. Despite our growing understanding of disease mechanisms in epilepsy, the current standard of care still relies on therapeutics seeking to address broad symptoms such as traditional anti-seizure medication (ASM) that globally suppress neuronal excitability through multiple mechanisms, including blocking of voltage-gated ion channels (e.g. carbamazepine), reduction of excitatory neurotransmission (e.g. topiramate) or augmentation of inhibitory neurotransmission (e.g. vigabatrin)^6,7^. However, these drugs can be associated with psycho-behavioral, cognitive and sleep adverse events^8^, underscoring the need for novel therapeutic compounds with improved properties and efficacy in treating specific genetic disorders.

One of the most prevalent among the known monogenic epilepsy disorders is Tuberous Sclerosis Complex (TSC), with an incidence of 1 in 6,000-10,000 newborns annually, and affecting approximately 2 million people worldwide^9,10^. TSC results from loss-of-function mutations in either the *TSC1* or *TSC2* genes, encoding for the proteins Hamartin and Tuberin, respectively, and which together form the TSC1-TSC2 complex^11^. Loss of function in this protein complex leads to hyperactivation of mTOR and associated pathways^11^. TSC patients develop benign tumors centrally and systemically in different tissues, with seizures and neurocognitive impairment as key clinical symptoms which can initiate in childhood and persist throughout adulthood^11^. Given the associated clinical manifestations and biological mechanism, therapeutic interventions have focused on traditional ASM such as early intervention with vigabatrin to reduce the risk of infantile spasms and drug-resistant epilepsy or mTOR inhibitors such as Everolimus that seek to target the underlying pathological mechanism^11^. However, not all seizures are addressed with the current approaches nor are the critical neurocognitive impacts such as intellectual disability and autism spectrum disorders^12,13^. In addition, small molecule ‘rapalogs’ (derivatives of the tool compound Rapamycin, a potent mTOR inhibitor and immunosuppressant) do target the underlying disease biology but lack the optimal drug-like properties required for maximum effectiveness and impact immune function, which can limit their clinical use^14,15^.

Given the continued need for improved therapeutic molecules and mechanisms, new drug discovery approaches are being developed that focus on scalable human cell-based models to better capture human physiological and pathological conditions in the context of patient-specific genetic backgrounds and disease-relevant cell types^16^. Several approaches have leveraged human induced pluripotent stem (iPS) cell-derived neuronal models of TSC to explore disease phenotypes at the cellular level and have identified changes in functional, morphological and transcriptional features, many of which could be rescued with Rapamycin^17–23^. These efforts represent a critical advance for the TSC field, demonstrating the initial proof-of-concept for the utility of a phenotypic-based approach using human cell-based models. Despite this significant progress, existing studies to date have been limited to the use of heterogeneous neural and neuronal cultures and low-power phenotypic or molecular studies focused on non-isogenic cell lines without validation in TSC patient-derived reagents or demonstration of *de novo* therapeutic screening^17–23^. Functional characterization in previous studies has focused on multi-electrode arrays (MEAs) which lack sufficient spatial resolution, or on Ca^2+^ imaging measurements which lack the temporal resolution to extract single action potential information. These studies^17–23^ have also generally utilized the tool compound Rapamycin to demonstrate cellular rescue and have not explored other rapalogs (including those used in the clinic) or other ASM. Lastly, no approach to date has demonstrated sufficient robustness and throughput of a phenotypic assay in a human neuronal model of TSC to drive a discovery screening campaign.

Here, we sought to create a functional platform for phenotypic CNS drug discovery applied to TSC-based epilepsy. To this end, we established diverse, scalable CRISPR/Cas9-isogenic and TSC patient-derived cellular models combined with high-throughput, single-cell resolved measurements of neuronal excitability to enable phenotype discovery across high-dimensional data sets. First, we used CRISPR/Cas9 tools to generate isogenic TSC iPS cell lines through the *knockout* of the *TSC2* gene in two independent genetic backgrounds, with multiple clones generated in each case. In parallel, we collected blood samples from TSC patients and related healthy controls and used these samples to generate iPS cell lines and neuronal reagents for downstream validation assays. These complementary cellular models allowed us to confidently identify phenotypes associated with TSC disease biology. Next, we developed an optogenetic assay system that allows for all-optical stimulation and recording of neuronal activity in a 96-well plate format with single-cell and single action potential resolution and sufficient throughput for small molecule screening^24,25^. We demonstrated the discovery of a multi-parameter electrophysiological disease phenotype along with RNA-seq based molecular signatures and changes to neuronal morphology, all of which are consistent with previous findings but provide deeper functional insights and substantially greater resolution at a scale appropriate for drug discovery. We then validated our TSC model by characterizing the extent of pharmacological phenotypic rescue by diverse mTOR modulators, including the clinical compound Everolimus. Lastly, we demonstrated the scale and robustness of the developed approach by conducting a phenotypic screen with ∼30,000 small molecule compounds in human iPS cell-derived neurons, which led to the identification novel chemical scaffolds that rescued the multi-parameter functional TSC disease phenotype. Notably, hits from the screen may act via different mechanisms than direct inhibition of the mTOR pathway. This strategy provided both molecular starting points for therapeutic development as well as a framework for phenotype discovery and drug screening in TSC-based epilepsy and other CNS disease states.

## RESULTS

### Generation of TSC human neuronal model and morphometric phenotype

To establish a human neuronal model of Tuberous Sclerosis Complex, we targeted the disruption of the *TSC2* gene, which negatively regulates the mTOR signaling pathway (**Figure 1A**). We used CRISPR/Cas9 gene targeting tools on two human iPS cell lines (11a and 20b) derived from neurologically healthy donors^26^ and generated a collection of *TSC2^+/+^*, *TSC2^−/−^* and *TSC2^+/−^* iPS cell clones (**Figure 1B**). Predicted *TSC2* genotype for each iPS cell clone, initially assessed by PCR and Sanger DNA sequencing, was confirmed with immunoblotting assays to determine TSC2 protein levels (**Figure 1C**). A subset of *TSC2^−/−^* iPS cell clones (each representing a different disruption in a different region of the *TSC2* gene) and *TSC2^+/+^* control cell lines (see **methods** for details) for each of the two genetic backgrounds was selected for further characterization. Isogenic *TSC2^+/+^* control iPS cells lines were generated from either the starting (Cas9-untreated) parental cell line or from an iPS cell clone in which *TSC2* editing did not occur despite treatment with Cas9 and gRNA reagents. Given the high efficiency of CRISPR/Cas9-mediated gene disruption, a single *TSC2^+/−^* heterozygous clone (∼50% TSC2 protein levels) for the 11a genetic background was identified after screening >150 iPS cell clones (**Figure 1C**).

**Figure 1.**
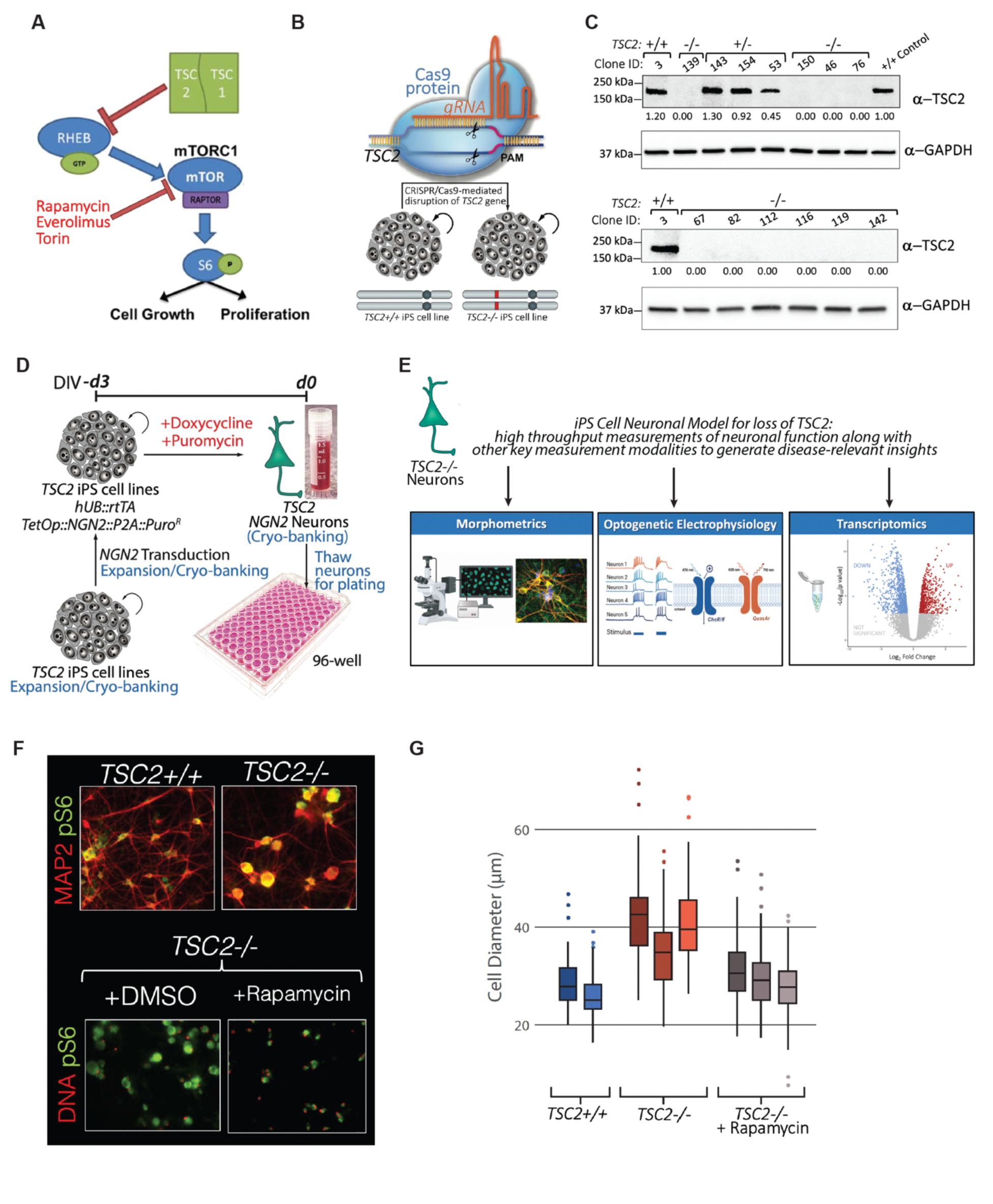
Generation of TSC human iPS cell neuronal reagents and validation of mTOR pathway-associated morphometric and pS6 phenotypes. **(A)** mTOR pathway: TSC1/TSC2 proteins negatively regulate the mammalian target of rapamycin (mTOR) complex 1 (mTORC1) signaling, a major regulator of cell growth and proliferation. **(B)** CRISPR/Cas9 was used on two control (*TSC2^+/+^*) iPS cell lines (11a and 20b) to generate a collection of *TSC2^+/+^*, *TSC2^−/−^* and *TSC2^+/−^* iPS cell clones. **(C)** Predicted *TSC2* genotype based on PCR Sanger Sequencing and TSC2 protein levels were confirmed using immunoblotting assays. **(D)** Cortical excitatory neurons (“NGN2” neurons) were produced from the different CRISPR/Cas9-edited isogenic *TSC2* clones using a transcriptional programming approach driven by overexpression of the proneuronal transcription factor NEUROG2 (NGN2). (**E**) High content imaging characterization of *TSC2^+/+^* vs *TSC2^−/−^* NGN2 neurons showed an increase in cell soma size and upregulation of pS6 kinase in *TSC2^−/−^* neurons, which could be reversed by chronic Rapamycin treatment, validating the disruption of mTOR signaling in these cells. **(F)** Immunocytochemistry of *TSC2^+/+^* and *TSC2^−/−^* neurons (detected with MAP2 antibody) showing upregulation of the mTOR effector pS6 in TSC2−/− neurons, which can be corrected with Rapamycin treatment. (**G**) Quantification of morphometric (cell size) phenotype and rescue with mTOR modulator Rapamycin

Having generated a collection of isogenic human iPS cell lines with different *TSC2* genotypes, we next differentiated them into cortical excitatory neurons using a transcriptional programming approach^24,27^ mediated by overexpression of the proneuronal transcription factor Neurogenin 2 (encoded by *NEUROG2* also known as *NGN2*) (**Figure 1D**). For each *TSC2* iPS cell line, we cryopreserved medium scale (∼20-100 x 10^6^ differentiated cells) production batches of these “NGN2 neurons” which we could then thaw, culture and characterize using morphometrics, optogenetic-based measurements of neuronal function, and transcriptomics (**Figure 1E**).

When cultured for 30 days, we detected an increase in cell soma size and upregulation of pS6 kinase in *TSC2^−/−^* neurons relative to *TSC2^+/+^* neurons, both of which could be rescued by chronic treatment with the mTOR inhibitor Rapamycin (**Figure 1F,G**). These phenotypes are well established readouts of disruption of TSC2 function and subsequent upregulation mTOR signaling^17,22,28^, and are consistent with the generation of a TSC cellular model in the human differentiated NGN2 neurons.

### Functional phenotypes in *TSC2^−/−^* neurons evaluated by optical electrophysiology

We interrogated the functional effects of TSC2 loss at the level of individual neurons using all-optical electrophysiology measurements. These assays were comprised of the blue light-activated channelrhodopsin CheRiff and the red light-emitting engineered voltage sensor QuasAr^29,30^ (**Figure 2A**). Lentiviral constructs expressing these two optical physiology components were delivered into NGN2 neurons of different *TSC2* genotypes co-cultured with primary mouse glial cells for 31 days, and measurements were collected using Quiver’s proprietary *Firefly* instrument^25^ (**Figure 2B**). Optical recordings were carried out in the presence of synaptic blockers to assess the functional intrinsic excitability firing pattern of each individual neuron (**Figure 2C**).

**Figure 2.**
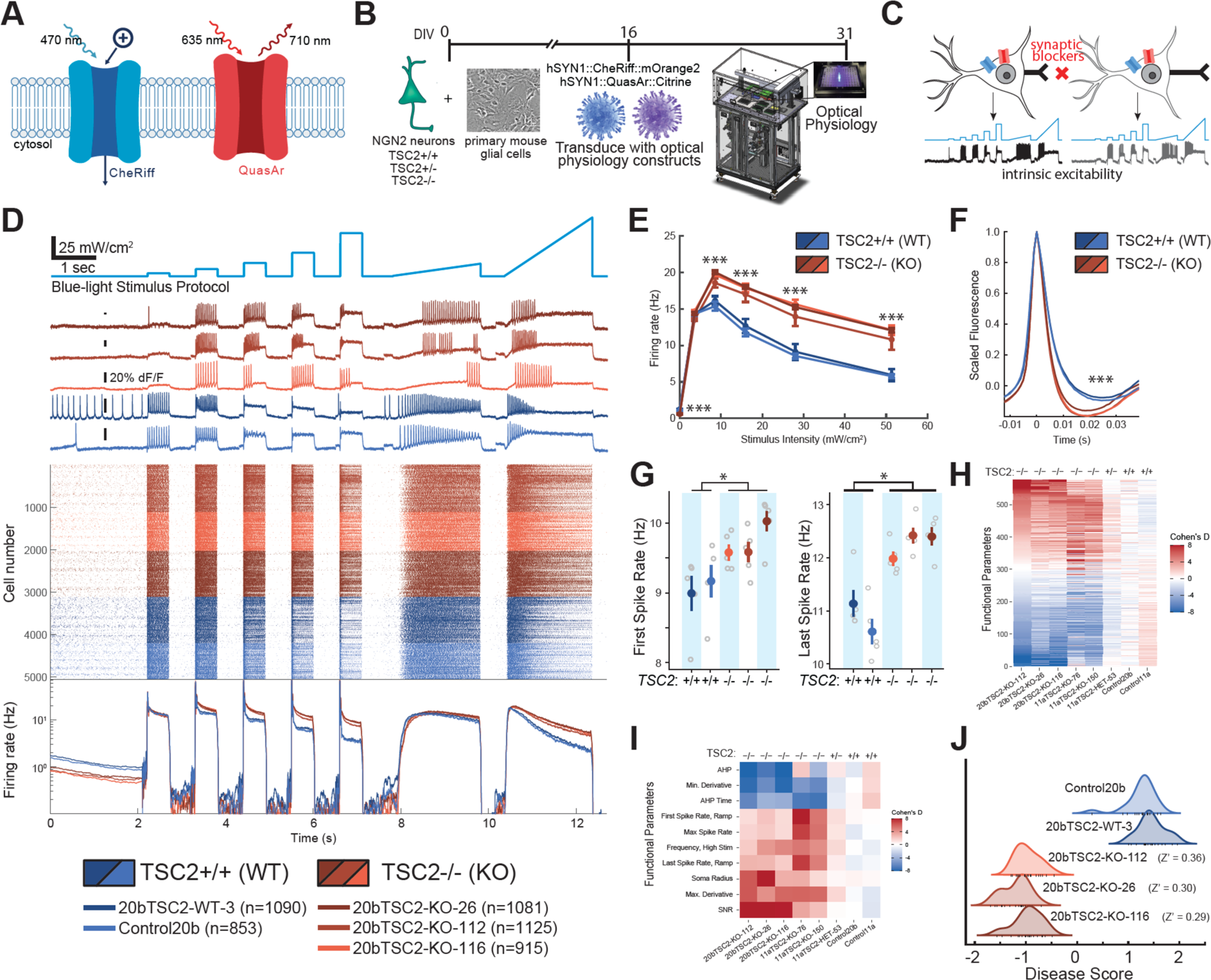
All-optical physiology platform reveals multiparametric intrinsic excitability phenotype in human iPS cell-derived neurons upon loss of TSC2 function. **(A)** Optical physiology platform is comprised of two genetically-encoded components, the optogenetic actuator CheRiff (a channelrhodopsin which stimulates neuronal activity in response to blue light) and the voltage reporter QuasAr (an archaerhodopsin which changes fluorescence emission in response to changes in membrane potential). **(B)** Assay timeline: human iPS cell-derived NGN2 neurons of different *TSC2* isogenic genotypes were plated and co-cultured with primary mouse glia for 31 days. CheRiff and QuasAr genetic constructs were delivered to all neurons via lentiviral transduction at day *in vitro* (DIV) 16, approximately two weeks prior to optical physiology imaging at DIV31. Functional measurements were collected using Quiver’s *Firefly* instrument. **(C)** Optical measurements were carried out in the presence of synaptic blockers to assess neuronal intrinsic excitability. **(D)** Functional characterization of one set of *TSC2^+/+^* and *TSC2^−/−^* isogenic neuronal reagents representing measurements of spike trains from >5,000 neurons and showing differential response to blue light stimulation in *TSC2* WT vs. *TSC2* KO genotypes. Top panel shows a fluorescence trace for a single exemplary neuron from each *TSC2* cell line; center panel shows a raster plot of action potentials from each the 5,064 neurons measured, with each cell represented by a row and each dot representing an individual action potential; bottom panel shows firing rate integral curve computed over all neurons in each group, on a log_10_ scale. **(E-G)** *TSC2* KO cells show statistically significant differences in spike timing and spike shape features when compared to *TSC2* WT neurons, including **(E)** differences in spike trains elicited by step stimuli (Frequency vs. Intensity, mean + SEM over wells), **(F)** differences in mean spike waveform (mean spike waveform + SEM over individual neurons), and **(G)** differences in spike trains elicited by ramp stimuli (spike rates at beginning and end of ramp stimulation, mean + SEM, individual data points are average values per well). **(H)** Heat map of the effect size and direction (*TSC2* KO vs. WT, signed Cohen’s D) for 577 features computed from optical physiology traces for the for the different *TSC2* iPSC clones compared to their corresponding isogenic *TSC2^+/+^* control. Features were sorted based on the TSC2-KO-112 vs. TSC2-WT-3 comparison (column 1), and this sorting was maintained to compare the multiple cell lines from the two genetic backgrounds (20b and 11a). **(I)** Heatmap with similar construction to **H**, but only considering a focused, diverse subset of phenotypic features. **(J)** Linear discriminant analysis (LDA) revealed that TSC2-KO-112 and TSC2-WT-3 have a larger phenotypic difference than other *TSC2* genotypic comparisons.

To determine whether loss of TSC2 function leads to altered intrinsic excitability, we measured the electrophysiological activity of NGN2 neurons derived from 5 different isogenic iPS cell lines, 3 *TSC2^−/−^*(*TSC2 knockout* or “KO”) and 2 *TSC2^+/+^* (TSC2 wild-type or “WT”) in the 20b^26^ genetic background. Functional activity was measured in response to patterned blue light stimulation designed to probe different neuronal response characteristics (weak/strong stimuli, steps/ramps, etc.) and data was collected from a total of 5,064 individual spiking neurons (**Figure 2D**). *TSC2^−/−^* neurons had lower activity than *TSC2^+/+^* neurons in the absence of optogenic stimulation but were hyperexcitable under high-intensity stimulation (**Figure 2D**). Similar results were observed for *TSC2* clones in the 11a^26^ genetic background (3 independent 11a CRISPR/Cas9-edited *TSC2^−/−^* clones vs. 2 isogenic *TSC2^+/+^* clones) (**Supplementary Figure 2**).

We identified individual neurons in each movie based on covarying pixel intensities (see **methods** for details) and then computed hundreds of functional features that describe dynamic patterns of action potential firing and action potential waveform shape in response to the blue light-mediated stimulus protocol used in the assay. *TSC2*^−/−^ and *TSC2*^+/+^ neurons exhibited statistically significant differences across 520 of the 577 functional parameters measured in these assays (two-way ANOVA, p<0.05 after false discover rate correction, see **methods** for details), indicating broad differences in neuronal behavior between the two groups (**Figure 2E-G, Supplemental Table 1**). Action potential frequency was higher in *TSC2^−/−^* neurons compared to *TSC2^+/+^* neurons under high-intensity stimulation, but lower at rest (**Figure 2D**, **2E**). The average shape of action potentials was also significantly altered by loss of TSC2, with spikes in *TSC2^−/−^* neurons showing narrower and deeper afterhyperpolarizations with steeper rising and falling slopes (**Figure 2F**). We also detected altered firing patterns in response to ramped stimulation in *TSC2^−/−^* neurons, with higher action potential frequency than *TSC2^+/+^* controls at both the onset and termination of firing (**Figure 2G**). In order to confirm that the functional phenotypes detected were not idiosyncratic to the 20b^26^ iPS cell genetic background, we also compared all 577 features across the second set of *TSC2* isogenic clones in the 11a^26^ iPS cell genetic background. For each feature of neuronal behavior, an average was taken over all individual neurons in a single well and signed Cohen’s D effect sizes were computed between groups of wells for each *TSC2* mutant genotype line and its corresponding *TSC2*^+/+^ control (TSC2-WT-3 for 20b cell lines, TSC2-WT-33 for 11a cell lines). The functional phenotype of *TSC2^−/−^* neurons was statistically significantly similar across multiple iPS cell clones in the two different genetic backgrounds (**Figure 2H**; Spearman rank correlation, p < 0.001). Note that when the starting (parental) *TSC2^+/+^* control cell lines (Control11a and Control20b) were compared to *TSC2^+/+^* clonal iPS cell lines (20b-TSC2-WT-3 and 11a-TSC2-WT-33), most of the features of these neurons showed effect sizes close to zero (**Figure 2H**). Interestingly, disruption of a single *TSC2* allele (*TSC2^+/−^*) in the 11a-HET-53 cell line resulted in effects that were smaller in magnitude but directionally similar for most features, showing a gene dosage effect on alterations of intrinsic excitability (**Figure 2H, Supplementary Figure 2**).

Finally, using a subset of statistically significant features chosen to represent key elements of neuronal behavior (**Figure 2I**), we applied a linear discriminant analysis (LDA)^31,32^ to compute a unidimensional ‘disease score’ representing an axis of phenotypic difference between all 20b *TSC2^−/−^* and *TSC2^+/+^* neurons (**Figure 2J**). Based on this analysis, the 20b TSC2-KO-112 cell line had the largest phenotypic difference (Z’=0.36) compared to the matched 20b TSC2-WT-3 control (**Figure 2J**). This 20b *TSC2^+/+^* (“WT-3” or “WT”) and *TSC2^−/−^* (“KO-112” or *“* KO”) isogenic pair was then selected for further studies described in the subsequent sections.

### Functional phenotype can be reversed by chronic, but not acute, administration of mTOR inhibitors

To further validate our TSC human neuronal model, we assessed disease-relevant pharmacology. Since loss of TSC2 function results in upregulation of mTOR activity^28^ (**Figure 1A**), we tested the effect of three pharmacological inhibitors of mTOR (Rapamycin, Torin-1, and the TSC clinical compound Everolimus) on the intrinsic excitability phenotype detected in *TSC2^−/−^* neurons. Each compound was tested on the 20b-TSC2-KO-112 neurons at two different concentrations and using three different treatment regimens (**Figure 3A**), ranging from early and chronic interventions [DIV10+24+31 (regimen A1)] to acute treatment [DIV31, 30-min incubation (regimen C1)]. Optical measurements of the neuronal behavior of *TSC2* KO neurons under the different conditions indicated that the longest chronic regimen (regimen A1) was required to induce the largest and most comprehensive reversal of phenotypic differences in features related to intrinsic excitability (**Figure 3B-C**). The shorter chronic regimen [DIV24+31 (regimen B1)] reversed some phenotypic features associated with firing rate but had no effect on spike shape features (**Figure 3C**), similar to acute treatment (regimen C1), which did not significantly alter the majority of phenotypic features. As early and chronic interventions (regimen A1) were necessary for phenotypic rescue by mTOR inhibitors, we chose to use a similar treatment regimen for downstream screening experiments.

**Figure 3:**
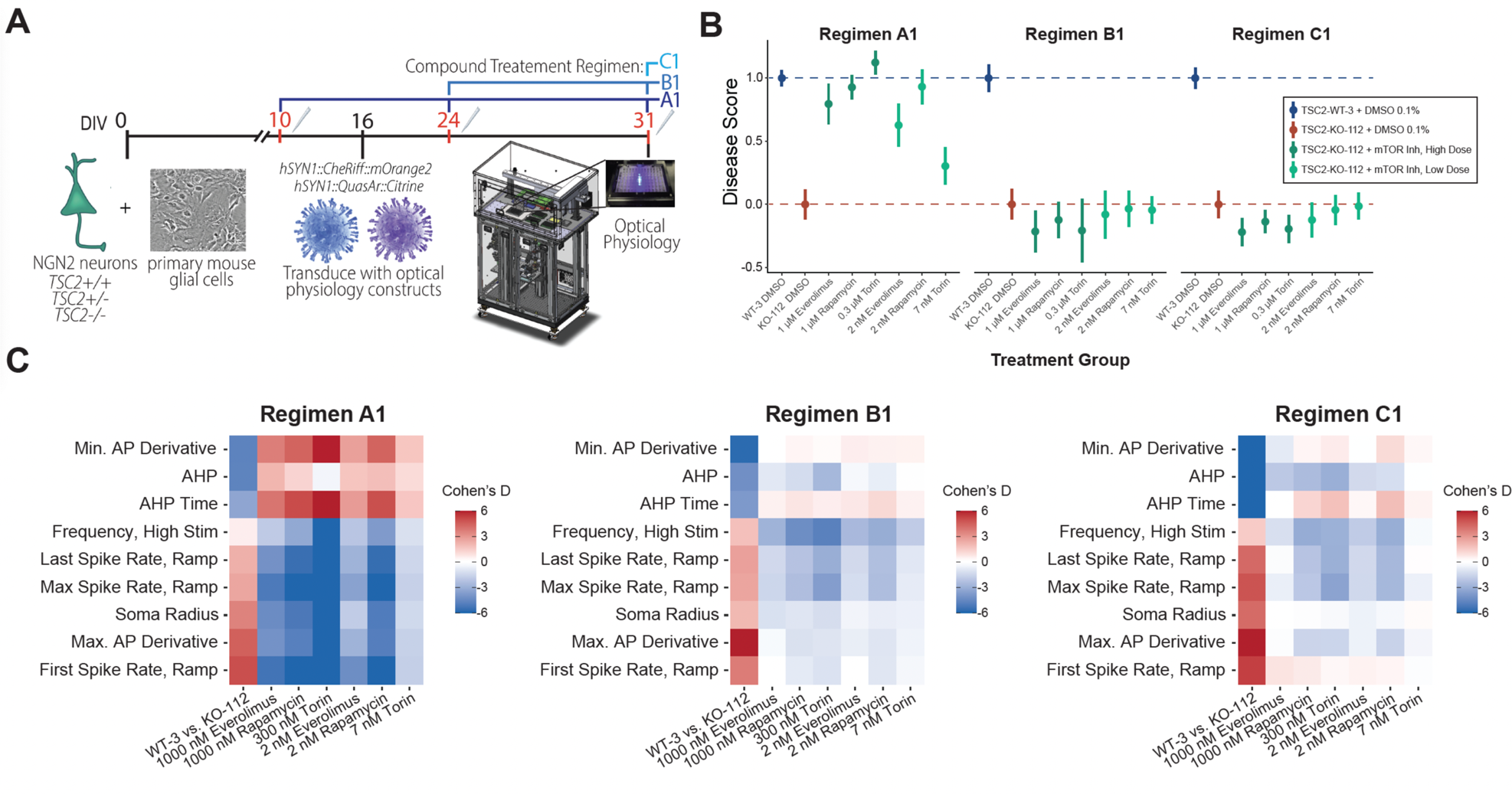
Chronic mTOR inhibition reverses functional excitability phenotype in *TSC2^−/−^* human neurons. **(A)** Assay timeline for cell culture, compound interventions (treatment regimens A1, B1 and C1), and optical physiology measurements. **(B-C)** Recovery of functional phenotype was tested using mTOR inhibitors at different doses and treatment regimens, revealing that long chronic dosing (Regimen A1) is necessary to see broad recovery of the functional phenotype, though partial recovery of some features is seen with shorter dosing regimens. **(B)** Functional phenotype rescue was quantified using a normalized LDA disease score, with assay wells for each condition normalized so that TSC2-KO-112 vehicle controls have a mean value of 0 while TSC2-WT-3 vehicle controls have a mean value of 1. **(C)** Magnitude and diversity of phenotypic rescue for each treatment regimen was plotted using a similar effect size heatmap used in Figure 2I, with the first column representing phenotypic differences between vehicle controls and remaining columns showing effect of three different mTOR modulators (at two different concentrations) on TSC2-KO-112 neurons. Experimental conditions showing a color opposite (in the Cohen’s D scale) to the phenotypic difference indicate functional phenotype reversal.

### Identification and rescue of molecular signatures in *TSC2^−/−^* human neurons

In order to interrogate molecular changes associated with the loss of TSC2 function in our human neuronal model, we used RNA-sequencing in two independent experiments to profile the transcriptomes of *TSC2* KO and *TSC2* WT iPS cell-derived NGN2 neurons co-cultured with mouse glial cells (**Figure 4A**). In the first experiment (“Exp 1”), cells were harvested for RNA-seq after 31 days in culture without additional experimental manipulations. In the second experimental round (“Exp 2”), neurons were treated with either 0.1% vehicle control (DMSO) or one of three mTOR pathway inhibitors (Everolimus, Rapamycin, and Torin-1) using three different treatment regimens before harvesting RNA at DIV31 (**Figure 4A**). Treatment regimens ranged from chronic and early interventions (administration at DIV10 and DIV24, regimen A2) to relatively “acute” (single administration 24 hours prior to harvesting RNA, regimen C2). When we compared genes that were significantly differentially expressed between the *TSC2* KO and *TSC2* WTgenotypes in untreated (Exp 1) or 0.1% DMSO-treated (Exp 2) neurons, we identified 7,080 genes that were consistently up-regulated or down-regulated in both experiments (**Figure 4B**). We focused our downstream analysis on these “high-confidence” genes with robust differential expression across two independent experiments.

**Figure 4.**
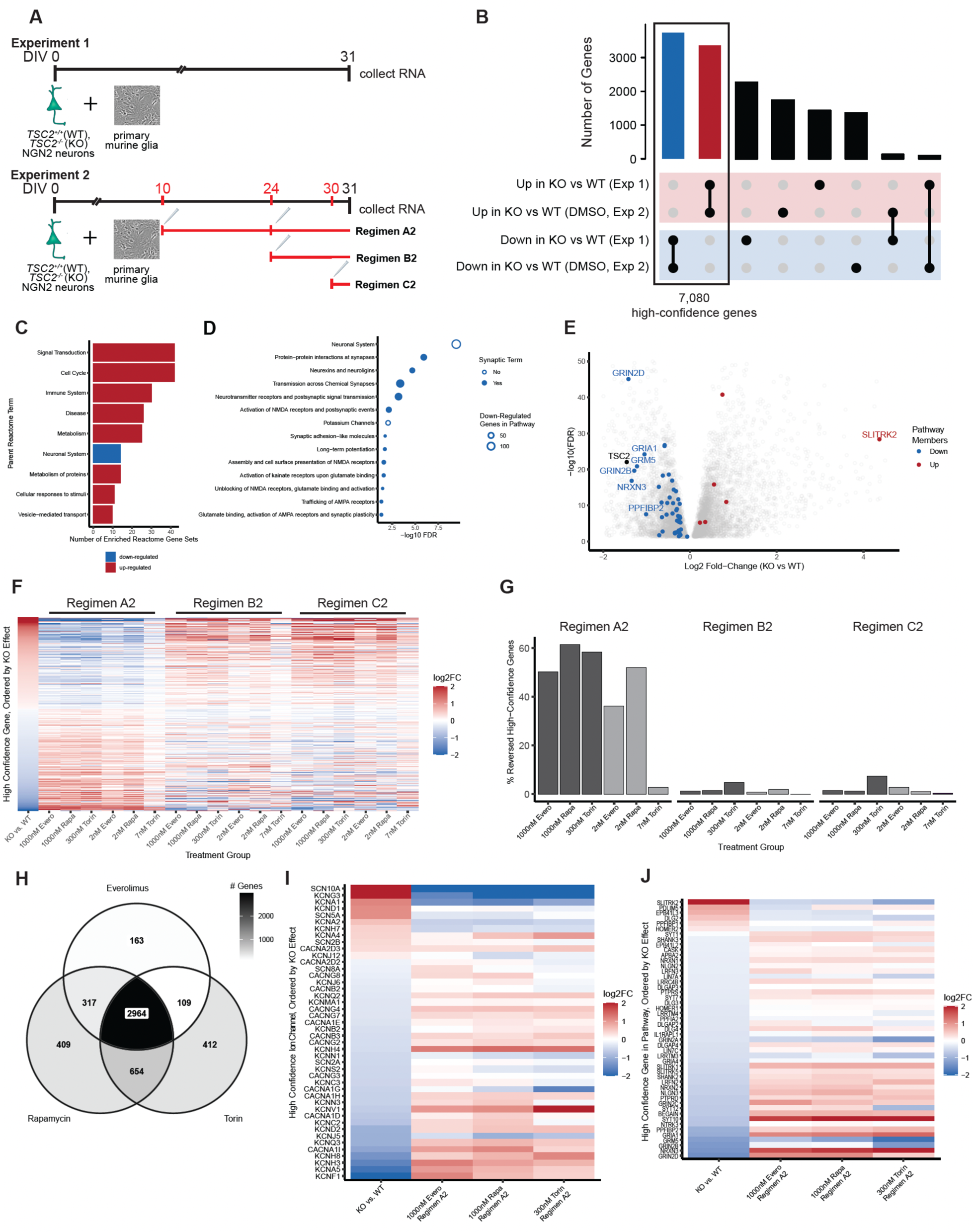
Transcriptomic profiling of *TSC2^−/−^* NGN2 neurons. (**A**) Timeline of two RNA sequencing experiments profiling *TSC2^−/−^* (TSC2-KO-112 or “KO”) and *TSC2^+/+^* (TSC2-WT-3 or “WT”) iPS cell-derived NGN2 neurons co-cultured with rodent glial cells. In Experiment 2, neurons were treated with 0.1% DMSO, Everolimus (“Evero”, 2 nM or 1000 nM), Rapamycin (“Rapa”, 2 nM or 1000 nM), or Torin (7 nM or 300 nM) using one of three treatment regimens. (**B**) UpSet plot of genes that are significantly differentially expressed between untreated (Exp. 1) or 0.1% DMSO-treated (Exp. 2) *TSC2* KO neurons and *TSC2* WT neurons. 7,080 “high-confidence” genes (red and blue bars) are shared. (**C**) Most frequent reactome pathway types enriched in high confidence differentially expressed genes. Top-level terms with ≥10 significantly enriched pathways are shown. (**D**) Neuronal systems reactome pathways significantly enriched in high-confidence down-regulated genes. (**E**) Volcano plot of 7,080 high-confidence genes in Exp. 2. Members of the “Protein-protein interactions at synapses” reactome pathway [R-HSA-6794362] are highlighted in red or blue. (**F**) Effect of drug treatment on high-confidence genes in Exp. 2. with genes sorted by the effect of *TSC2 knockout* in 0.1% DMSO-treated neurons (far left column). All other columns compare drug-treated *TSC2* KO neurons to 0.1% DMSO-treated *TSC2* KO neurons in that treatment regimen. Log_2_ fold-change values >2 or <-2 have been flattened to 2 or −2. (**G**) Percentage of the 7,080 high-confidence genes whose expression is significantly reversed by compound treatment in *TSC2* KO neurons across compound and treatment regimens. (**H**) Overlap between genes that are significantly reversed by high dose regimen A2 treatments of Everolimus, Rapamycin, and Torin in *TSC2* KO neurons. (**I**) Effects of *TSC2 knockout* (far left column) or high dose regimen A2 treatments of mTOR modulators on *TSC2* KO expression of high-confidence genes encoding ion channels. Log_2_ fold-change values >2 or <-2 have been flattened to 2 or −2. (**J**) Effects of high dose regimen A2 treatments on high-confidence genes in the “Protein-protein interactions at synapses” reactome pathway [R-HSA-6794362]. Log_2_ fold-change values >2 or <-2 have been flattened to 2 or −2.

We next investigated the function of this subset of differentially expressed genes using gene ontology analyses. High-confidence genes that are up-regulated in *TSC2* KO neurons include members of reactome pathways involved in signal transduction, regulation of the cell cycle, immune response, disease, and metabolism, while high-confidence down-regulated genes include members of pathways important for neuronal function and communication (**Figure 4C**). Many of these high-confidence down-regulated genes are members of pathways that govern synaptic behavior (**Figure 4D**). The most down-regulated members of the “Protein-protein interactions at synapses” pathway [R-I-6794362] mediate glutamate signaling, including genes encoding the NMDA receptor subunits GRIN2D and GRIN2B, AMPA receptor subunit GRIA1, and metabotropic glutamate receptor GRM5 (**Figure 4E**). These results are consistent with previously reported deficits in synaptic behavior associated with mutations in *TSC2* genes^28^. Furthermore, our own preliminary optogenetic-based characterization of synaptic transmission in cultured human *TSC2* KO neurons showed reduction in the amplitude of synaptic events in these cells and that chronic treatment with mTOR inhibitors led to partial recovery of this synaptic phenotype (**Supplementary Figure 3**).

Finally, we evaluated whether treatment with mTOR pathway inhibitors (Rapamycin, Torin, and Everolimus) could rescue the molecular signatures caused by loss of TSC2. We found that early and chronic treatment (regimen A2) with high doses of all three compounds could reverse the direction of a large fraction of the high-confidence genes in *TSC2* KO neurons, with 1000 nM doses of Rapamycin in regimen A2 reversing the direction of over 60% of the altered genes (**Figure 4F-G**). The other two treatment regimens (regimens B2 and C2) largely failed to reverse the direction of expression of the high-confidence genes, which suggests that, similar to the rescue of electrophysiological deficits (**Figure 3C**), chronic treatment with mTOR compounds is necessary to restore many of the molecular consequences of TSC2 loss. Consistent with the shared target pathway of the three compounds tested, the genes reversed by the high dose regimen A2 treatments of the three compounds largely overlapped (**Figure 4H**). A number of ion channels (**Figure 4I**) and synaptic function genes (**Figure 4J**) are dysregulated in *TSC2* KO neurons, and many of these transcriptional changes are partially reversed by treatment with mTOR modulators. These molecular changes may contribute to the functional phenotypic features and rescue of intrinsic excitability (**Figures 2**, **3**) and synaptic behavior (**Supplementary Figure 3**) observed in our all-optical physiology experiments.

### Validation of functional phenotype in patient-derived cell lines

To provide further validation of the intrinsic excitability phenotype associated with the loss of TSC2 in the CRISPR/Cas9 isogenic disease model, we also carried out functional measurements in TSC patient-derived neurons. To this end, we collected blood samples from three TSC patients with confirmed genetic loss of *TSC2* along with samples from their unaffected relatives (one of the parents of the TSC patient with no reported genetic change in *TSC2*). We generated iPS cell lines from all six donors and differentiated them into NGN2 neurons. TSC patient (*TSC2^+/−^*) and healthy control (*TSC2^+/+^*) neurons were plated alongside the CRISPR/Cas9-isogenic genotypes 20b-TSC2-KO-112 and 20b-TSC2-WT-3 for all-optical electrophysiology characterization. We observed a similar pattern of changes in intrinsic excitability in TSC patient vs. control cell lines (n=11,394 neurons) and the isogenic CRISPR/Cas9 model (n=3,349 neurons). Action potential frequency is reduced in patient lines relative to matched controls at rest but elevated under high-intensity stimulation (raster integrals in **Figure 5A**, **5B**, **2D**; frequency-intensity (F/I) curve in **Figure 5C**, **5F**, **2E**). A similar pattern of changes between the TSC patient-derived and CRISPR/Cas9 models was also observed in the firing activity of the neurons during the ramp stimulus (**Figure 5E**, **5H**, **2G**). Lastly, average action potential waveform shape in TSC patient-derived neurons showed a similar significant reduction (p < 0.01) in spike width and similar significant increase (p < 0.01) in after hyper-polarization (**Figure 5D**) as those detected in the isogenic model (**Figure 2F**, **5G**). Together, these results support the validity of the CRISPR/Cas9 *TSC2^−/−^* isogenic reagents in modeling TSC-associated functional phenotypes given the strong concordance with the phenotype observed in multiple TSC patient-derived models.

**Figure 5:**
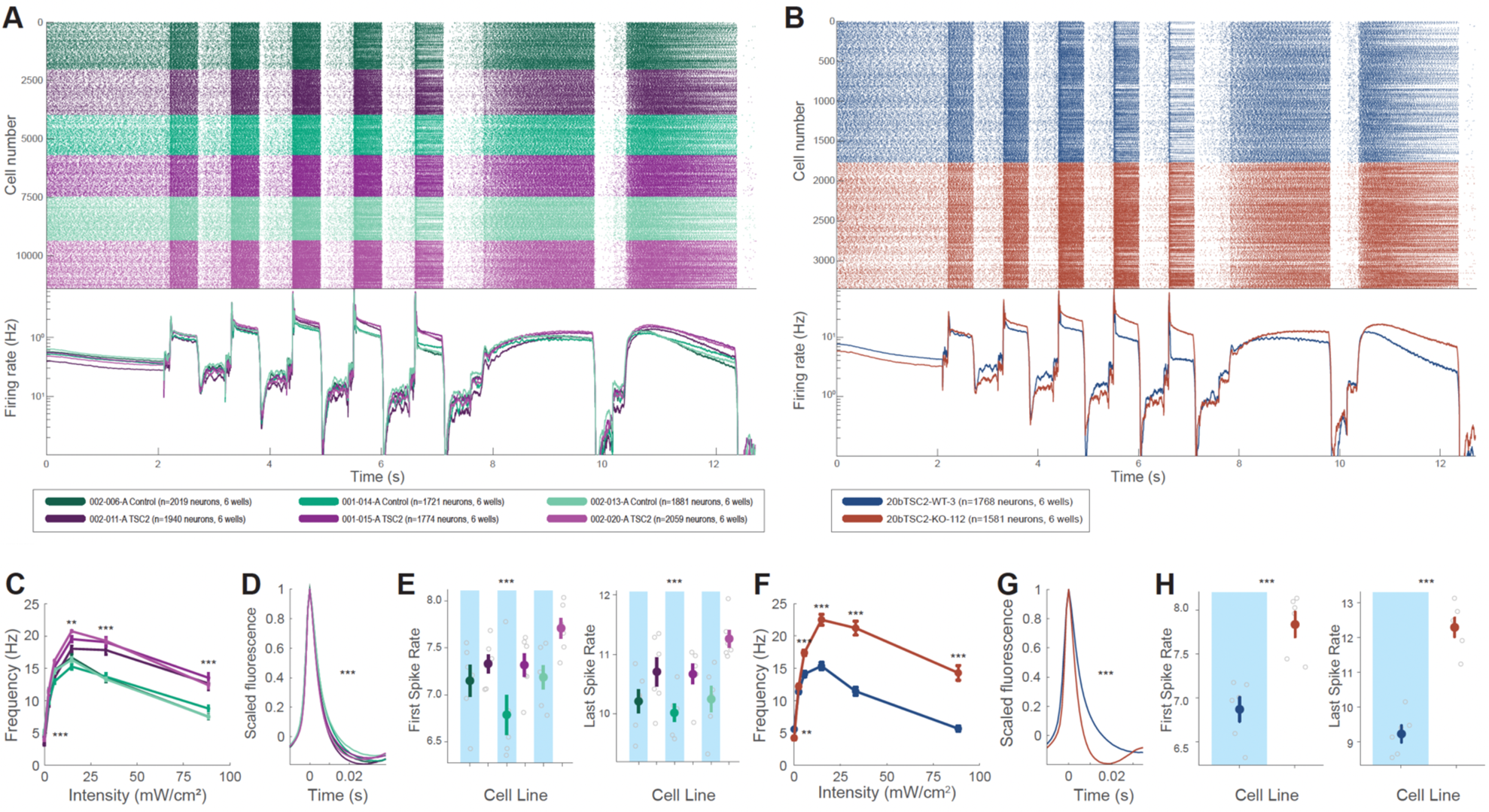
Functional phenotype in TSC patient-derived NGN2 neurons matches phenotype detected in CRISPR/Cas9 TSC2-KO iPS cell-derived neurons. Similar phenotypic differences are seen in firing rate integral (**A** vs **B**); F/I curves (**C** vs **F**); action potential waveform shape (**D** vs **G**); and response to ramped light stimulus (**E** vs **H**).

Interestingly, while our isogenic model is a full knockout of *TSC2* expression, TSC patient-derived cells are *TSC2^+/−^* and showed several of the same functional features associated with complete loss of TSC2 function. Characterization of a single CRISPR/Cas9 11a *TSC2^+/−^* clone (11a-HET-53) indicated that a subset of functional features identified in the *TSC2^−/−^* neurons are significantly affected and to a lesser degree by the single *TSC2* allele disruption (**Figure 2H**, and **Supplementary Figure 2**), suggesting that at least in this 11a iPS cell control genetic background disruption of both *TSC2* alleles may be necessary for neurons to manifest a stronger intrinsic excitability phenotype that matches those identified in patients with TSC. Characterization of additional isogenic *TSC2^+/−^* clones may be needed to fully understand the contribution of mono and bi-allelic *TSC2* mutations, as well as genetic background to the neuronal functional phenotype. Nevertheless, given the strong overlap in phenotypic feature space from the *TSC2*^−/−^ and those observed in multiple TSC patient lines, as well as the robust phenotype window, we proceeded with compound screening activities using the isogenic model system.

### Multiparametric phenotypic screen identifies >400 confirmed hits from a ∼30,000 small molecule compound library

Having established a robust functional phenotype and a pharmacological rescue strategy in human *TSC2^−/−^* neurons, we next sought to conduct a high throughput phenotypic screening assay to drive the identification of novel chemical scaffolds modulating the human neuronal model of TSC. To this end, we used a 31-day culture assay with 20b-TSC2-KO-112 human neurons treated with small molecules at 3 different time points (DIV10+24+31) prior to functional data acquisition (**Figure 6A**). Using this assay, we reliably measured a phenotypic window with an effect size that was compatible with high-throughput screening (Z’>0.4, **Figure 6B**) using the LDA disease score developed in the initial phenotypic characterization (**Figure 2I-J)**. Pharmacological tests indicated that this disease score could be used to measure recovery of the phenotype by reference compounds (**Figure 6B**). For the primary high throughout functional screen, we measured the effect on the *TSC2* KO phenotype of each of 29,250 small molecule compounds dosed at 5 μM (3 interventions, n=1 well per compound) on >300 96-well plates, which yielded the identification and parameterization of 7,043,437 individual neurons (**Figure 6C**, **6D**). On each day of functional imaging and data acquisition, we measured the behavior of a “sentinel” plate, which contained only positive (20b-TSC2-WT-3 neurons) and negative (20b-TSC2-KO-112) phenotypic controls as well as positive rescue pharmacology (mTOR modulator Torin-1 and potassium channel modulator XE-991). Any imaging round for which the sentinel plate Z’ was < 0.4 was discarded and repeated; therefore, as a result, all sentinel plate Z’ values considered for hit selection were > 0.4 (**Figure 6E)**.

**Figure 6.**
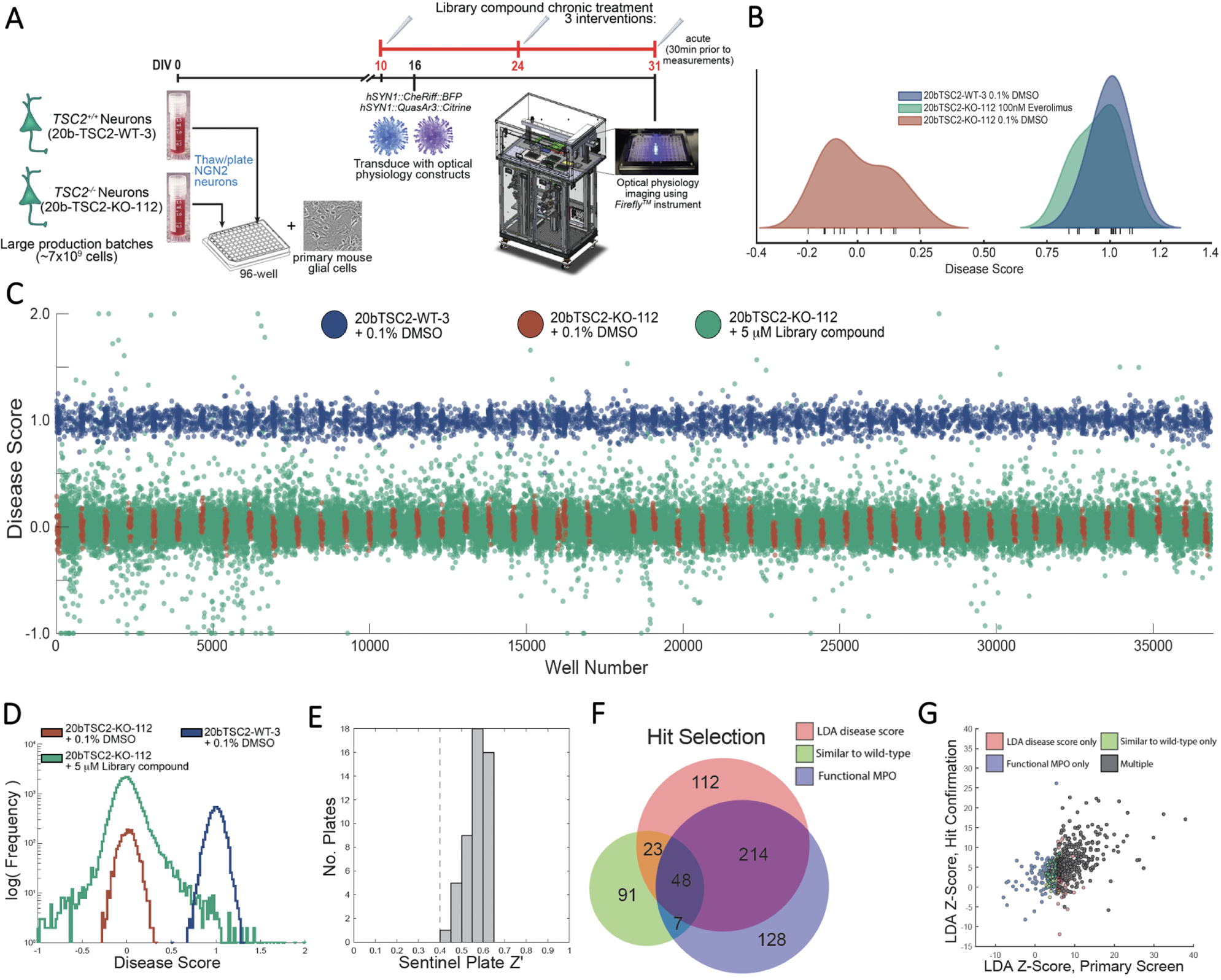
Completion of a ∼30,000 small molecule compound phenotypic screen and subsequent hit confirmation enabled the identification of molecules that reversed the functional phenotype of *TSC2* KO neurons. **(A)** Assay timeline: NGN2 neurons for 2 isogenic *TSC2* genotypes [*TSC2^+/+^*(20-TSC2-WT-3) and *TSC2^−/−^* (20b-TSC2-KO-112) were produced in large batches (billions of cryopreservable differentiated neurons) to support high throughput screening and downstream activities. A 21-day compound treatment regimen (with 3 interventions) was chosen for the primary screen. **(B)** LDA (linear discriminant analysis) disease scores were computed and normalized to illustrate a screening window (*Z’* = 0.41) between the TSC2-WT-3 and TSC2-KO-112 neuronal lines. Treatment of TSC2-KO-112 neurons with 100 nM Everolimus for 21 days modulates the disease scores of TSC2-KO-112 cells to overlap with TSC2-WT-3 scores. **(C)** Disease score metric for each library compound-treated well from the primary phenotypic screen was plotted in the order of optical physiology imaging. Each day of data acquisition included a sentinel plate which contained positive (TSC2-WT-3 neurons, in blue) and negative (TSC2-KO-112 neurons, in brown) control wells, followed by several screening plates, which contained positive control wells and library compound wells (in green), for a total of 336 screening plates and 49 sentinel plates, representing n=5,000 total positive control wells, n=1,960 total negative control wells, n=29,250 library compound testing wells. Disease score values outside the plot limits [-1,2] have been clipped to appear at the limits. **(D)** Histogram of disease score metric from panel (C), plotted on log scale and showing enrichment for active compounds in the tails of distribution. **(E)** Histogram of *Z’* values for sentinel plates throughout the primary screen. **(F)** A total of 623 hits were identified using three algorithms. This Venn diagram shows the amount of overlap among the three approaches, each of which identified many unique compounds. **(G)** For hit confirmation, disease score was normalized as a z-score of the negative controls. Scatter plot shows a significant correlation between primary screen and hit confirmation values for all 623 hits.

We devised three methods to identify hits in order to maximize their chemical and mechanistic diversity. First, all disease scores were transformed into a z-score scale relative to the negative controls on the corresponding sentinel plate, and the top 400 compounds ranked by LDA disease score were selected (LDA score method) (**Figure 6F**). These 400 compounds exhibited normalized LDA values between 0.29 and 2.94 (on the scale used in **Figure 6C**), corresponding to z-score values between 5.73 and 41.4 standard deviations over control. Second, all of the individual features included in the LDA disease score were converted to z-scores by the same method, then those z-scores were summed to produce a ‘functional MPO (multi-parametric optimization)’ score, and the top 400 compounds ranked by this functional MPO method were selected (**Figure 6F**). Finally, any compounds that recovered the phenotype such that every feature included in the LDA disease score was within 3 standard deviations of the mean of the TSC2-WT controls (similar to wild-type method) were also included (**Figure 6F**). Each approach added a significant population of unique compounds to the pool of 623 hits identified, with 48 compounds overlapping all three methods (**Figure 6F**). These findings suggested that a diversity of potential independent mechanisms may underlie the functional rescue by the hit compounds identified in our phenotypic screen.

To confirm the hits from the primary screen, we retested the 623 compounds in duplicate (n=2 wells) and at two different concentrations (5 μM and 1 μM) using the same treatment regimen (DIV10+24+31). LDA scores in the primary screen and hit confirmation screen were significantly correlated (Pearson’s rho=0.48, p < 0.001, **Figure 6G**). Hits in this secondary screen were classified based the z-transformed disease score (>3σ from controls). Based on this metric, the overall hit confirmation rate across categories of hits was 70% (with 14% also active at 1 μM). For the 400 hits that were initially selected based on primary disease score, the rate was 77%. In total, we identified 434 compounds from the primary screen that were confirmed in the hit confirmation screen.

### Most hit compounds are unlikely to act via direct mTORC1 modulation

Following hit confirmation, we performed additional analyses to further understand whether the hits identified in our phenotypic screen are likely acting via modulation of the mTOR pathway or through alternative mechanisms. *TSC2^−/−^* neurons exhibit increased levels of pS6 expression that can be reduced by chronic treatment with mTORC1 pathway inhibitors (**Figure 1F**, **Figure 7A**). We found that chronic treatment with the majority of the 623 hit compounds at 1 µM or 5 µM does not modify this increased pS6 immunoreactivity phenotype, suggesting novel mechanisms of action (**Figure 7A**). Of the 2,487 total wells treated with either 1 µM or 5 µM of a hit compound from the primary screen, only 16 have a normalized pS6 expression value <0.5. In contrast, all 84 wells treated with the mTOR inhibitor Torin-1 have values below 0.5. XE-991, a potassium channel blocker that rescues much of the functional phenotype in *TSC2* KO neurons, notably does not modify the pS6 expression phenotype (**Figure 7A**).

**Figure 7.**
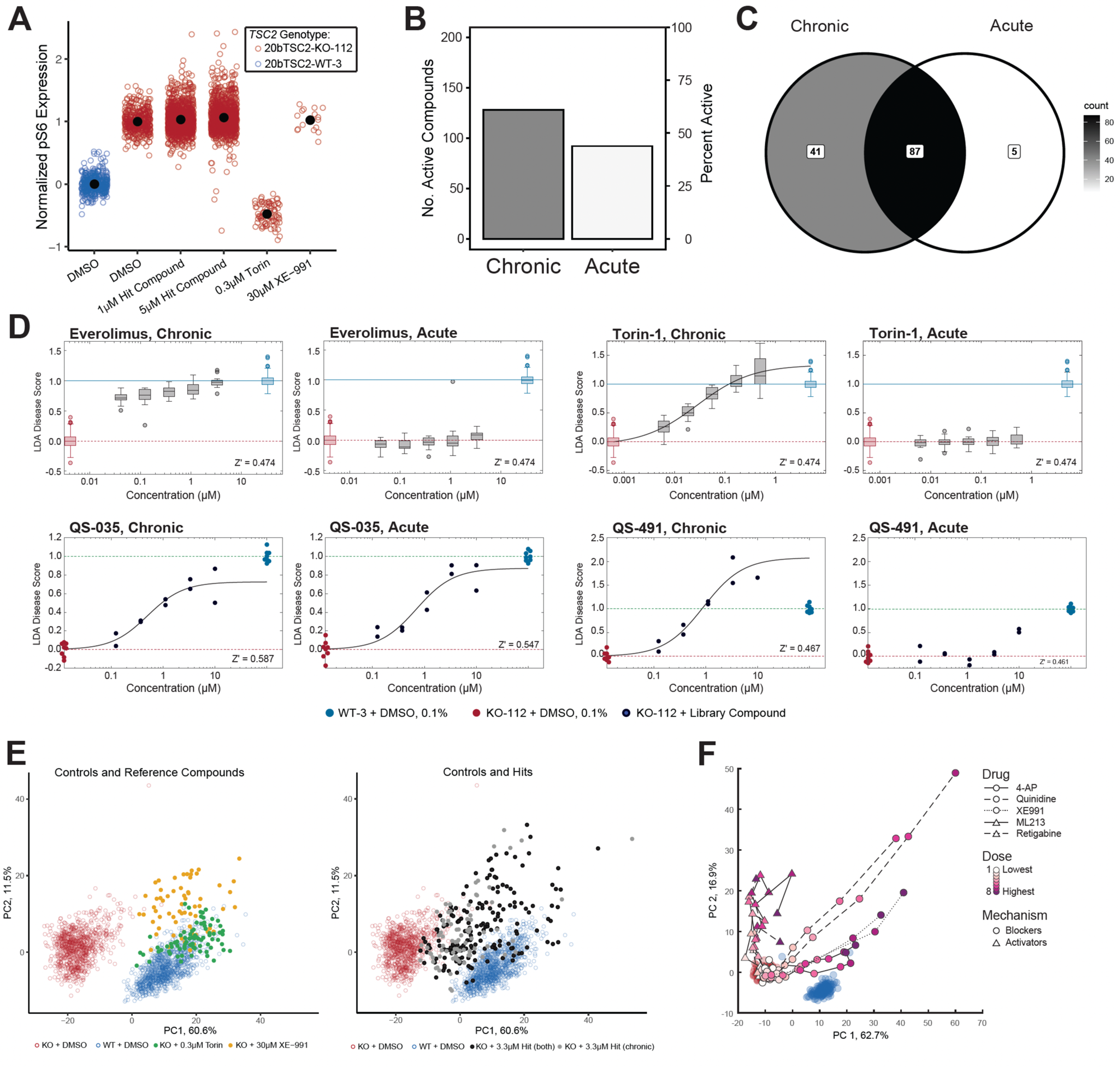
Characterization of hit compounds. **(A)** Normalized pS6 expression in *TSC2^+/+^* neurons treated with DMSO (n= 464 replicate wells) and *TSC2^−/−^* neurons treated with DMSO (n= 432 wells), 1 μM hit compound (n=1246 wells, 2 replicates per compound for 623 hit compounds), 5 μM hit compound (n=1241 wells, generally 2 replicates per compound), 0.3 μM Torin-1 (n=84 wells), or 30 μM XE-991 (n=16 wells). **(B)** Number of 210 top hit compounds active (LDA>0.3) at 3.33 μM in a chronic (DIV10+24+31) or acute (30-90min) treatment regimen. **(C)** Overlap between active groups in B. (**D)** Example concentration-response curves for two mTOR modulators (Everolimus and Torin-1) and two top hit compounds (QS-035 and QS-491). **(E)** Principal component analysis of functional behavior in a chronic treatment regimen for *TSC2* KO neurons treated with 0.3 μM Torin-1 or 30 μM XE-991 (left panel) or 3.33 μM library compound colored by activity category (right panel). DMSO-treated *TSC2* KO and *TSC2* WT controls are shown in both panels. (**F)** Principal component analysis of functional behavior in a chronic treatment regimen for *TSC2* KO neurons treated with potassium channel blockers or activators vs. DMSO-treated KO and WT controls.

Compounds that do not act via modulation of mTORC1 may not require the long chronic dosing regimen used during the phenotypic screen to rescue the *TSC2* KO neuronal excitability behavior. To investigate this hypothesis, we next characterized a focused set of 210 confirmed hits in 5-point dose response using both the chronic screening regimen (DIV10+24+31) and an acute dosing regimen (compound addition 30-90 minutes) prior to optical physiology measurements (treatment regimens A1 and C1, respectively, from **Figure 3A**). These 210 focal compounds were chosen from the pool of 623 hit candidates based on confirmed phenotypic recovery, the absence of activity in the pS6 assay, potency, and the attractiveness of the chemical scaffold for future development. Compounds with an LDA disease score >0.3 at the 3.33 µM dose were classified as active in this experiment. We found that 61.0% of the hit compounds are active when administered chronically and 43.8% of hit compounds are active when administered acutely (**Figure 7B**). Most of these compounds are active under both treatment regimens (**Figure 7C**), further suggesting mechanisms of action distinct from mTOR modulators, which are only active chronically, as shown by their dose-response curves (Everolimus, Torin-1, **Figure 7D**). While the EC_50_ for chronic administration of Torin-1 was estimated to be 28.3 nM, because of the range of concentrations we tested in this particular assay, we cannot estimate the EC_50_ for Everolimus, but it was certainly less than 30 nM. In addition to these two mTOR modulators, we present example concentration response curves for two hit compounds: compound QS-035, which can restore the functional phenotype after either chronic or acute treatment, and compound QS-491, which requires chronic treatment for phenotypic rescue (**Figure 7D**). Both of these confirmed hits represent promising molecules for future development, with QS-035, given its acute phenotype rescue effects, likely acting through novel mechanisms distinct from direct mTORC inhibition.

Finally, we investigated whether we could identify clusters of hits with similar properties based on their overall functional profile. Hits in our study were selected based on the LDA score parameters that best separate the functional behavior of *TSC2^−/−^* and *TSC2^+/+^* neurons (**Figure 2I-J**, **Figure 6B**, **6F**), but these parameters represent only a subset of the hundreds of functional features we collected during the screen. In a principal component analysis of the functional behavior of *TSC2* KO neurons chronically treated with 0.3 µM Torin-1, 30 µM XE-991, or 3.3 µM hit compounds and DMSO-treated *TSC2* WT and *TSC2* KO controls, we found that the functional behavior of neurons treated with a known potassium channel blocker and a known mTOR modulator can be distinguished along the second principal component (**Figure 7E**, left panel). However, hit compounds classified as only chronically active or both chronically and acutely active (**Figure 7C**) are not well-separated in the first two principal components of this landscape (**Figure 7E**, right panel). Given the variance in the locations of XE-991 and Torin-1-treated neurons and range of potencies seen in the hit compounds, additional replicates and additional doses could improve separation of compounds in this landscape. Moreover, the inclusion of additional reference compounds with divergent mechanisms could provide insights into mechanisms of action. For instance, in a similar analysis performed on a separate functional electrophysiology dataset, we found that potassium channel blockers and potassium channel activators have distinct functional profiles when applied to *TSC2* KO neurons and that multiple compounds from the same class have similar profiles (**Figure 7F**). We think that a landscape constructed with our functional platform using a large, diverse library of drugs with known mechanisms of action could facilitate future studies and accelerate target deconvolution efforts.

## DISCUSSION

Tuberous Sclerosis Complex (TSC), a rare genetic disorder caused by loss of function mutations in the mTOR pathway genes *TSC1* or *TSC2*, results in CNS symptoms which include intractable seizures and cognitive deficits^9,10^. Appropriate human neuronal models that can recapitulate disease phenotypes are critical for the discovery and development of novel and improved therapeutics to address these neurological symptoms^23,33^. In this study, we developed a human neuronal model of TSC which both recapitulates and builds upon morphological, molecular, and electrophysiological phenotypes previously described in other models of TSC2 deficiency^23,33^. The scale and depth of our optical physiology characterization represents a significant advancement in understanding the functional impact of TSC2 loss in disease-relevant human neuronal models. We further validated the functional phenotypes of our human cell-based model by characterizing both CRISPR/Cas9-generated and TSC patient-derived cellular reagents from multiple genetic backgrounds. We then used our all-optical electrophysiology platform to conduct a highthroughput phenotypic screen at an unprecedented scale for functional measurements in human iPS cell-derived neurons, ultimately identifying >400 confirmed hit compounds that rescued the neuronal excitability phenotype associated with loss of TSC2.

A successful phenotypic screen using human iPS cell-derived neurons requires a high-quality, disease-relevant cellular model. We generated a collection of TSC-relevant human neuronal reagents by applying transcriptional programming methods to both TSC patient-derived and CRISPR/Cas9-edited isogenic iPS cell lines. Similar to a recent study^22^, we found that using a direct and rapid approach mediated by overexpression of the NEUROG2 transcription factor bypassed some of the proliferation phenotypes reported in neural *TSC2^−/−^* progenitors when small molecule-mediated neuronal differentiation methods were used^17,19^. We did not observe deficits in neuronal production efficiency by the different *TSC2^−/−^* iPS cell lines (**Supp Figure 1**), which is an important factor for minimizing potential confounding variables in downstream CNS disease modeling assays^34^. Similar to previous reports^17,22^, we also observed that human iPS cell-derived neurons with loss of TSC2 showed increase soma size and upregulation of pS6 immunoreactivity (**Figure 1F-G**). As these two readouts are well-established phenotypes of mTOR signaling hyperactivation downstream of TSC2 deficiency^10,11^, these findings served as an initial validation of our cellular TSC model. The validity of this TSC neuronal model was further supported by the rescue of many functional and molecular disease signatures in *TSC2^−/−^* neurons using tool (Rapamycin and Torin-1) and clinical (Everolimus) mTOR-modulating compounds (**Figure 3**).

The utility of a functional phenotypic assay is based on its ability to reflect key properties of the disease and to facilitate the identification of candidate therapeutics or molecular targets. In TSC, patients suffer seizures and neurocognitive deficits, which can occur independent of the presence of tubers in the CNS^9,10^, suggesting that loss of TSC1/2 can have direct effects on neuronal function. By using a proprietary all-optical electrophysiology platform with single-cell and single-action potential resolution^24^, we identified a robust excitability phenotype in cultured human neurons caused by loss of TSC2. This neuronal excitability phenotype cannot be broadly classified as ‘hypoexcitability’ or ‘hyperexcitability’, but rather includes functional parameters associated with both. In *TSC2^−/−^* NGN2 neurons, we observed reduced action potential firing when there was no stimulus applied, no difference at low stimulus intensity, and increased firing in response to moderate and high stimulus intensity (**Figure 2D**, **2E**, **2I**). The observed reduced neuronal firing in *TSC2^−/−^* neurons during the spontaneous period is consistent with previous reports describing the loss of TSC function^17,20,35,36^, but published F-I (frequency-current) curves are inconsistent among the different studies. Some reports show reduced firing in neurons lacking TSC signaling during stimulation^17,35^, while others are consistent with our observation of elevated firing rates (**Figures 2E**, **5C**, **5F**) even at the stimulus intensity that elicits peak firing rate^36^. Some of these discrepancies in neuronal behavior can be attributed to the differentiated cell type and resting membrane potential properties (e.g. Purkinje cells^20^ and mixed excitatory/inhibitory culture^17^ in previous reports vs. NGN2 excitatory neurons in this study). The changes we detected in spike shape properties of *TSC2^−/−^* neurons, which included narrower action potentials (APs) with deeper afterhyperpolarization (AHP), could partially explain the higher firing rates during high stimulus intensity as narrower APs with deeper AHP may be expected to rebound faster. Importantly, the functional differences between *TSC2* KO and *TSC2* WT neurons were broadly consistent across CRISPR/Cas9-isogenic *TSC2^−/−^* neurons from two genetic backgrounds and the TSC patient-derived *TSC2^+/−^* reagents (**Figure 5A, B**). A single *TSC2^+/−^* heterozygous isogenic iPS cell line displayed a functional phenotype that was similar to that of the *TSC2^−/−^* neurons, albeit with smaller amplitude changes (**Figure 2I**, **Supplementary Figure 2**). While patients are heterozygous for loss of *TSC2*, the concordance of the disease signatures in *TSC2*^−/−^ isogenic lines with those of the patient-derived neurons supported the use of a *TSC2*^−/−^ isogenic line as a sensitized disease model for our phenotypic screen.

In addition to functional phenotyping, our transcriptomics analyses of *TSC2^+/+^* and *TSC2^−/−^* human iPS cell-derived neurons detected >7000 molecular signatures associated with loss of TSC2 (**Figure 4B**). Transcripts that were significantly downregulated are enriched in genes related to neuronal function (**Figure 4C, I**). Inspection of ion channel transcripts that are differentially expressed in *TSC2^−/−^* versus *TSC2^+/+^* neurons suggests candidate genes to explain observed electrophysiological changes. Increased expression of sodium channel genes is consistent with increased action potential upstroke velocity, and upregulation of voltage-gated potassium genes is consistent with an observed increased repolarization rate and an increased afterhyperpolarization depth. Downregulation of a number of voltage-gated potassium and voltage-gated calcium channels could be mechanistically associated with the spike timing and spike initiation changes observed in *TSC2^−/−^* neurons, but additional experiments would be needed to connect expression of specific gene signatures with specific electrophysiological changes. Additionally, consistent with mutations in *TSC* genes leading to alterations in synaptic transmission^28^, we found a large number of synaptic genes to be dysregulated in *TSC2^−/−^* NGN2 neurons (**Figure 4C**), which provides a means to investigate possible molecular mechanisms underlying synaptic deficits that we detected in preliminary assays (**Supplementary Figure 3**). These synaptic gene expression signatures in *TSC2^−/−^* neurons included glutamate-gated ion channel subunits, voltage-gated calcium channels, and voltage-gated potassium channel subunits that are associated with epileptic encephalopathies, providing a possible mechanistic link to the seizures observed in TSC patients.

We believe the transcriptional, morphological, and functional alterations in *TSC2^−/−^* neurons that we report in this study can represent a link between molecular and biochemical changes following reduction in TSC2 activity and network level changes that can manifest in patient clinical phenotypes. The use of a human neuronal model of TSC in a prolonged 30-day culture, allowed us to screen using multiple chronic and acute intervention points (**Figure 6A**) that were informed by the molecular and functional rescue assays using mTOR modulators, including the clinical compound Everolimus (**Figure 3C**, **4G**). Of note, while other groups have developed and characterized functional neuronal models for TSC, including the use of Ca^2+^, MEA and patch-clamp measurements^23^, our proprietary all optical electrophysiology platform made it possible to undertake a high-throughput functional phenotypic screen using a library of ∼30,000 small molecule compounds representing both high chemical diversity and CNS drug-like properties. Over the duration of this study, including phenotype discovery, assay optimization, primary phenotypic screen, hit confirmation, and follow-up characterization of hits, we extracted hundreds of spike shape and spiking timing parameters from a total of 20,336,348 individual neurons and over 1.653 billion action potentials. To our knowledge, this is the first report of a phenotypic screen of this scale and complexity using electrophysiological data in cultured human stem cell-derived neurons.

The confirmed hit compounds identified in our functional phenotypic screen provide a key starting point for additional screening and optimization steps, such as SAR (structure activity relationship) characterization with compound analogues, early ADME (Absorption, Distribution, Metabolism, and Excretion) studies, exploration of medicinal chemistry, validation in other human neuronal types, *in vivo* rescue of TSC genetic mouse models, and target deconvolution efforts. The elucidation of target and mechanism of action represents a fundamental challenge in phenotypic drug discovery^37^. While the primary screen described here was performed using a treatment regimen appropriate for the identification of known therapeutic compounds, many of the hit compounds appear to differ from a TSC therapeutic compound (Everolimus) in their mechanism of action. Most of the confirmed hit compounds do not rescue elevated pS6 levels in *TSC2^−/−^* neurons and, interestingly, many of the hit compounds rescue the functional disease phenotype under an acute treatment regimen (**Figure 7A**, **7B**). A subset of these hits may act on channels or on more proximal mediators of neuronal excitability, similar to XE-991, a potent and selective K_v_7 blocker that modifies the disease phenotype but is separable from the mTOR inhibitor Torin-1 in the high-dimensional phenotypic space (**Figure 7E**). This type of analysis suggests that a reference space developed using the phenotypes of human neurons treated with a library of compounds with known mechanisms of action could facilitate target deconvolution.

In sum, by establishing a robust cellular model of TSC, we completed an unprecedented multiparametric functional phenotypic screen of ∼30,000 small molecules in human iPS cell-derived neurons and identified >400 confirmed hits. These compounds appear to be distinct from current therapies and can be used to generate novel and differentiated molecules for TSC drug development. Importantly, our human functional platform can easily be extended to other genetic disorders of the nervous system with underlying neuronal excitability phenotypes. Finally, our characterization of compound treatment regimens and target mechanisms suggests that our approach may also provide a framework for the elucidation of novel target mechanisms for TSC and other CNS-based disorders.

## METHODS

### Derivation of human iPS cell lines from TSC patients

Under IRB-approved protocol and consent forms and in collaboration with Dr. Orrin Devinsky (NYU Langone and Saint Barnabas Institute of Neurology and Neurosurgery) we obtained peripheral blood samples (2–4 mL) from TSC patients with confirmed loss-of-function (premature stop codon) mutations in *TSC2* and from healthy relatives (one of the patient’s parents). Peripheral blood mononuclear cells (PBMCs) were then isolated by density gradient centrifugation with Ficoll. PBMCs were reprogrammed into iPS cells using transient overexpression of pluripotency-associated factors (OCT4, KLF4, and SOX2) via non-integrating Sendai virus particles (Cytotune 2.0 kit, Life Technologies). At least five clones per *TSC2^+/+^* and *TSC2^+/−^* donor were established into iPS cell lines. *TSC2* genotype was confirmed in the resulting iPS cell lines by PCR and Sanger DNA sequencing and lack of TSC2 protein expression in the TSC2 patient-derived lines was validated using immunoblotting assays.

### Production of *TSC2^+/+^*, *TSC2^+/−^* and *TSC2^−/−^* isogenic iPS cell lines using CRISPR/Cas9

Two control iPS cell lines, 11a and 20b^24,26^ were used to establish a collection of isogenic lines with different *TSC2* genotypes. These two iPS cell lines were initially derived from two neurologically healthy male donors and have been characterized in previous neuronal studies^24,26^. gRNAs targeting either Exon 11, Exon 17, or Exon 23 in the *TSC2* gene were cloned into a plasmid expressing Cas9 and a puromycin resistance gene. 11a and 20b iPS cell lines were transfected with the Cas9+gRNA expressing plasmid and after transient antibiotic selection, puromycin resistant colonies were individually expanded to establish multiple *TSC2^+/+^* and TSC2^−/−^ iPS cell clones in both 11a and 20b genetic backgrounds. A single 11a *TSC2^+/−^* clone was identified. TSC2 genotypes were determined using Sanger DNA sequencing of PCR amplicons spanning the *TSC2* gene region targeted by the respective *TSC2* gRNA.

### Culture of human iPS cell lines and neuronal production

All iPS cell lines (CRISPR/Cas9-edited isogenic clones and TSC patient-derived lines) were routinely expanded in culture and maintained using mTeSR1 medium (STEMCELL Technologies) and cell culture surfaces pre-coated with Matrigel (Corning) according to manufacturers’ recommendations. iPS cell lines were differentiated into cortical excitatory “NGN2” neurons using a transcriptional programming approach, as previously described^24,27^. Briefly, iPS cell lines were initially transduced with lentiviral particles expressing the reverse tetracycline transactivator (rtTA) and a tetracycline-responsive (TetOp) construct driving the expression of the proneuronal transcription factor Neurogenin-2 (NGN2) and a puromycin antibiotic resistance enzyme. These genetically modified iPS cell lines were then expanded in mTeSR1 medium for three to six passages prior to induction of NGN2 expression via doxycycline. For neuronal production, iPS cells were dissociated with Accutase (STEMCELL Technologies) according to manufacturer’s recommendations and plated at a density of 300,000 cells/cm^2^ using mTeSR1 medium supplemented with 10 μM Rock Inhibitor (Sigma) and 2 mg/mL doxycycline (Sigma) to initiate NGN2 overexpression. A day later, the medium was switched to a 1:1 DMEM/F-12: Neurobasal Medium (Thermo Fisher Scientific) supplemented with 1x GlutaMAX (Life Technologies), 1x non-essential amino acids (Life Technologies), 1x N2 (Gibco), 2 mg/mL doxycycline (Sigma), and 2 μg/mL puromycin (Sigma). Medium was exchanged daily with fresh medium for three additional days, which then resulted in a highly homogenous population of post-mitotic neurons, which were then dissociated with Accutase and cryopreserved at densities of 10-20 million cells per cryovial. For the primary phenotypic screen, a single large batch of 20b-TSC2-KO-112 (*TSC2^−/−^*) and 20b-TSC2-WT-3 (*TSC2^+/+^*) neurons was produced by scaling up the expansion of iPS cell lines modified with rtTA/NGN2 constructs to allow for the production and cryobanking of at least 7×10^9^ cells, sufficient to support the primary screening activities. To initiate neuronal assays, cryopreserved neurons were thawed and plated onto poly-D-lysine/laminin pre-coated 96-well culture plates (Ibidi^TM^) at a density of 60,000 cells/cm^2^ on day *in vitro* (DIV) 0. Human neuronal cultures were maintained for 30 days in Neurobasal A Medium (Life Technologies) supplemented with 1x GlutaMAX (Life Technologies), 1x non-essential amino acids (Life Technologies), 1x N2 (Gibco), 1x B-27, 10 ng/mL BDNF (R&D), and 10 ng/mL GDNF (R&D). To allow for neuronal maturation, the differentiated human cells were co-cultured with mouse primary glial cells which were obtained as previously described^38^ and seeded on DIV0 at a density of 40,000 cells/cm^2^ in each well.

### Immunocytochemistry assays

Immunocytochemistry (ICC) assays were carried out as previously reported^24,39^. Briefly, iPS cell-derived neuronal cultures were fixed using 4% paraformaldehyde diluted in 1 PBS from a 16% aqueous solution (Electron Microscopy Sciences) for 20 min and then permeabilized with 0.2% Triton X-100 (Sigma). Cell culture dishes were blocked with 10% donkey serum in 0.1% PBS-Tween for 1 h and then incubated with primary antibodies overnight at 4C. After five washes with 0.1% PBS-Tween, dishes were treated with secondary (Alexa-conjugated) antibodies for 1 h at room temperature. Cell nuclei (DNA) were stained with Hoechst 33342. Primary immunoreagents used in this study included antibodies detecting the cytoskeletal pan-neuronal markers MAP2 (microtubule associated protein 2) [Abcam; ab5392 (1:3,000) or Novus Biological; NB300-213 (1:3000)] and β-III TUBULIN [TUJ1, BioLegend; #801202 (1:1,000)], a human nuclear-specific antigen [(EMD Millipore; MAB1281 (1:1000)], as well as the mTOR effector phosphor S6 ribosomal protein (pS6 kinase) [Cell Signaling; #4858 (1:1000)]. Images were acquired as maximum-intensity Z-projections using the automated confocal microscope GE Healthcare IN Cell Analyzer 6000 with 20 (0.75NA) objective.

### Immunoblotting assays

For detection of protein levels using immunoblotting assays, protein concentrations were determined using BCA assay on a Nanodrop, and 10–20 mg of protein sample were loaded and separated using SDS-PAGE. After protein transfer onto membranes, these were incubated with primary antibodies overnight and corresponding HRP-conjugated secondary antibodies for 1–2 h prior to developing and measuring protein signal. Primary antibodies and dilution used were TSC2 (Cell Signaling Technology 1:1000 dilution) and GAPDH (Abcam ab9484, 1:2,000 dilution).

### Functional assays using all-optical physiology

Optical physiology assays were carried out as previously described^24^, with some minor modifications. Briefly, distinct lentiviral particles encoding the optogenetic actuator CheRiff and QuasAr were produced in 293T cells as previously reported^30,40^. Neuronal cultures were transduced around two weeks prior to functional imaging (on DIV16) with CheRiff and QuasAr constructs driven by the human *SYNAPSYN 1* (*hSYN1*) promoter to target expression of the optical physiology components to mature neurons. Transduction of neuronal cultures was carried out as described before^30^, with neuronal culture medium exchange 16–24 h after transduction to remove remaining lentivirus. Neuronal cultures were maintained until DIV30-31, when neuronal function was recorded using a custom built ultra-widefield fluorescence microscope^25,30^, the Firefly. Briefly, samples were illuminated with 1) ∼ 100 W/cm^2^ 635 nm laser excitation (Dilas) to monitor changes in membrane potential through changes in QuasAr fluorescence and 2) variable blue light intensity (0–110 mW/cm^2^, 470 nm LED, Luminus) to depolarize the neuronal membrane through CheRiff. Custom blue light stimulus protocols were used to probe different facets of the electrophysiological response, as described in the text. Imaging data were recorded on a Hamamatsu ORCA-Flash 4.0 sCMOS camera across a 4 mm x 0.5 mm field of view at a 1 kHz frame rate. Data acquisition was performed using custom control software written in MATLAB.

### Functional imaging movie analysis and feature extraction

Fluorescence time-series for every neuron were extracted from each movie using a PCA-ICA algorithm based on unique covariance of pixel intensities of each functional neuron to identify unique, statistically independent masks that correspond to each spiking neuron^29,41^. The time series for each neuron were obtained by computing the dot product of the mask with the movie tensor. Quality metrics were computed on the mask (size, radius, compactness, etc.) and time series (number of spikes, signal-to-noise ratio (SNR), etc.) and only neurons that passed pre-defined quality thresholds are retained for further feature extraction and analysis. Action potentials (APs) were detected using a kernel-based method with criteria to reject false positives. For each spike, several parameters were computed to describe the waveform shape: rise-time (in frames, from spike initiation to peak amplitude), width (in ms, at 20% of the AP height), depth of the afterhyperpolarization (as a fraction of spike height), time of peak afterhyperpolarization (in ms, from spike initiation), and the minimum and maximum of the first and second derivatives during the spike. For each epoch of the imaging protocol, an average shape parameter was computed to represent all spikes under that kind of stimulus (shape, intensity, duration). We also computed several parameters describing the spike train in each epoch of the imaging protocol, including frequency, latency to first spike, time to last spike, a measure of spike frequency adaptation, and the first, last, and maximum instantaneous frequencies between pairs of action potentials. Finally, to gain additional resolution, each epoch of the imaging protocol was divided up into time bins of varying length (50-200 ms) and spike rate and shape features were aggregated for spikes within those bins, for greater sensitivity to dynamic changes to different parts of the stimulus (onset, steady-state, post-stimulus, etc.). Note that because of these factors the exact number of features we computed depends on the stimulus protocol and changed throughout the results section as we optimized the stimulus protocol for screening. Because our optical physiology platform can produce thousands of voltage imaging movies per day, all movie analysis and feature extraction were conducted using cloud compute services. Shortly after collection, each movie was uploaded to S3 storage on Amazon Web Services (AWS; Amazon, Seattle, WA) and analyzed using our AWS’ elastic compute service. Results were stored on S3 and loaded into a MySQL database for further analysis.

### Statistics for functional phenotype identification

Statistical evaluations were carried out with well-level averages – for each feature and well measured we computed an average value over all neurons in that well. This has several advantages, including addressing missingness in our single-cell-level data and reducing the effect of intraclass correlations on downstream statistical analyses. Because we measured a large number of features of neuronal behavior from each experiment, we evaluated statistical significance for each feature independently and then adjusted for multiple comparisons using the false discovery rate (FDR) algorithm ^42^. Specifically, we conducted a two-way ANOVA for each feature, with the two dimensions being disease status (*TSC2* KO vs. *TSC2* WT) and cell line, and evaluated significance based on the FDR-adjusted p-value for the main effect of disease status. Where appropriate, we conducted post-hoc t-tests for each feature, correcting for multiple comparisons using Bonferroni’s method.

### Disease score estimation of functional phenotype

In order to carry out screening activities, we generated a reduced-dimensionality measure of the disease phenotype. For simplicity, we elected to use a linear discriminant analysis (LDA) to compute a weighted average of disease features. To estimate stable weights for the LDA model using limited data (dozens of wells per experiment relative to hundreds of possible features), we selected a representative subset of features to use in computing the LDA disease score. Features were selected based on several factors, including statistical significance, effect size, robustness over multiple experiments, feature importance reported by machine-learning classifiers, and relationship to previously published literature on physiological phenotypes associated with TSC2 knockout. During high-throughput screening, each day of imaging included a sentinel plate which consisted of phenotypic controls (health, disease) and reference compounds for quality control. The phenotypic controls on the sentinel plate were used to evaluate the LDA weights that are used to compute disease scores for all other wells, including library compounds, for each day of imaging.

### RNA-Sequencing

Human iPS cell-derived NGN2 neurons were cultured as described above. At DIV30 (experiment 1) or DIV 31 (experiment 2) cells were washed in PBS, incubated in TRIzol at room temperature for 10 minutes, and then pooled (3 wells/sample) and stored at −80°C prior to RNA extraction. In experiment 1, n=12 replicate samples/condition (TSC2 genotype) were collected. In experiment 2, n=4 samples/condition (drug, dose, and treatment regimen combination in one genetic background) for Everolimus, Rapamycin, or Torin-1-treated neurons and n=8 samples/condition for DMSO-treated neurons were collected. RNA extraction was performed using Qiagen RNeasy plus mini kits. For experiment 1, sequencing libraries were prepared using NEBnext Ultra RNA directional kit (#E7420) and 2×151 sequencing was performed using a NovaSeq 6000. For experiment 2, libraries were prepared using an Illumina TruSeq Stranded RNA Library Preparation Kit and 2×150 paired-end sequencing was performed using a HiSeq 4000. For both experiments, sequencing reads were filtered and trimmed using bbduk/BBMap (v38.31) to remove adapters (kmer: 20mers), low quality sequences (Phred<20), short reads (<31nt), and low complexity or homopolymer reads (Entropy filter: 0.01). Trimmed reads were pseudo-aligned to a human/mouse composite reference transcriptome (gencode human v29, mouse vM19, kmer size 31) and quantified using Salmon v0.11.3^43^. Differential expression analysis was performed in R (3.5) using Bioconductor packages (release 3.8). Human transcript counts were summarized to the gene level using tximport and scaled using library size and the average transcript length over samples^44^. Counts were TMM-normalized, log2 transformed, and quantile normalized^45^. Data from experiment 2 was additionally batch-corrected using combat^46^. Lowly expressed genes were removed (experiment 1: genes where <10 samples had at least 10 counts; experiment 2: genes where <4 samples had at least 30 counts). Differential expression between conditions was assessed for each experiment independently with limma using voom for variance estimation and sample weights^47,48^. These models included technical covariates for experiment 1 (glial cell batch, production batch, differentiation day).

*Identification and characterization of high-confidence genes*: 7,080 “high-confidence” genes were significantly differentially expressed in consistent directions in experiment 1 (TSC2^−/−^ KO-112 vs. TSC2^+/+^ WT-3) and experiment 2 (DMSO-treated TSC2^−/−^ KO-112 vs TSC2^+/+^ WT-3, all treatment regimens combined). A hypergeometric test was used to assess the enrichment of Reactome pathway members in the high-confidence genes relative to detected genes. The number of significantly enriched pathways was summed by the highest parent-level reactome terms represented in v3.7, Reactome database release 82 of the reactome pathway browser (https://reactome.org/PathwayBrowser/). Synaptic pathways were defined as “Protein-protein interactions at synapses [R-HSA-6794362]”, “Transmission across Chemical Synapses [R-HSA-112315]”, “Transmission across Electrical synapses [R-HSA-112307]”, or any downstream child of these three pathways. High-confidence ion channel genes were identified by intersecting the 7,080 gene symbols with the following lists of ion channel genes from the Human Gene Nomenclature Committee gene sets (https://www.genenames.org/data/genegroup/#!/group/292, downloaded 11/04/22): “Sodium voltage-gated channel alpha subunits”, “Sodium voltage-gated channel beta subunits”, “Sodium leak channels, non selective”, “Calcium channel auxiliary gamma subunits”, “Calcium voltage-gated channel alpha1 subunits”, “Calcium voltage-gated channel auxiliary alpha2delta subunits”, “Calcium voltage-gated channel auxiliary beta subunits”, “Potassium calcium-activated channels”, “Potassium voltage-gated channels”, “Potassium inwardly rectifying channel subfamily J”, “Potassium sodium-activated channel subfamily T”, and “Potassium two pore domain channel subfamily K”. Results were summarized and visualized in R (3.6) using tidyverse packages, ggplot2 (3.3), ComplexHeatmap (2.2), and ggVennDiagram (1.2.2). RNA sequencing raw and processed data files will be deposited in GEO.

### Small molecule library for phenotypic screening

We screened a ∼30,000 small molecule compound library, which was composed of two smaller libraries (∼15,000 compounds each) selected from two separate pharmaceutical compound library collections. Compounds in each library were selected based on structural diversity and CNS drug-like properties. Both ∼15,000-compound library sets were selected from the corresponding larger collections using the same selection criteria described below. The first ∼15,000-compound focused library was generated from a master library of ∼200,000 commercially available small molecules compounds. About half of the 200,000 compound library was selected to maximize diversity and explore chemical space. Most of the remainder compounds were chosen for good CNS-drug-like characteristics based on molecular properties. Extensive filters were applied during selection to remove liabilities such as reactive groups, chelators, auto-florescence, and aggregators, and to yield compounds amenable to eventual medicinal chemistry optimization. To select the 15,000 compound set from the master 200,000 library, compounds were sorted into 9000 structural groups by Extended Connectivity Fingerprint (ECFP) analysis^49,50^ with maximum dissimilarity = 0.65 and assigned a CNS Multiparameter Optimization (MPO) score^51^ based on the Pfizer MPO model with no modifications to the scoring system which predicts CNS drug-like properties. Compounds that were potentially reactive, aggregating, chelating, or auto-fluorescent were removed, and PAINS^52^, acids and zwitterion compounds were flagged for future review. ∼15,000 compounds were selected to maximize structural diversity with maximized MPO score within each structural group. ECFP analysis clustering indicated that this ∼15,000-compound library contained >94% of the structural diversity in the 200K master library; 87% have an MPO score ≥5. The same approach was used to select ∼15,000 compounds from a larger compound library provided by the pharmaceutical company UCB (srl, Braine l’Alleud, Belgium). The final library used in this study contained ∼30,000 compounds with less than 5% overlap between the individual libraries based on chemical similarity and complementarity, yielding a library that maximized structural diversity across the contributing compound collections.

### High throughput screening analysis, hit selection, and hit confirmation

We performed a phenotype screen on ∼30,000 small molecule compounds to identify compounds able to reverse the *TSC2* KO intrinsic excitability phenotype. Initial experiments were used to optimize assay conditions, including DMSO tolerance, compound treatment regimen and screening concentration. The 30,000 compound library was designed to maximize chemical diversity and CNS drug like properties for use in focused high throughput screens (HTS). The screen used NGN2 neurons from a batch of ∼7 billion neurons to ensure consistency across the campaign. Day-to-day variability was evaluated by including a sentinel plate containing alternating columns of *TSC2* KO and *TSC2* WT cells for each screening day. In cases when Z’ calculated from a sentinel plate did not match or exceed 0.4, all screening plates from those days were scheduled for retesting. The intrinsic excitability data generated for the HTS assay contains a rich set of descriptors of single cell neuronal function and the assay platform enabled collection of the large datasets needed to robustly identify functional disease phenotypes. We applied dimensionality reduction approaches to this large dataset to generate a small number of quantitative metrics to rank compounds. Linear Discriminant Analysis applied to the >500 features contained in the intrinsic excitability data set to generate a one-dimensional LDA disease score for all cells in a well, provided a means to assess phenotype rescue by test compounds. Each well in a screening plate contained ∼100-200 functional neurons; analysis of variations in disease score at the single well level demonstrated a screening window (Z’≥0.4) sufficient for enabling screening using a single well per compound. Using a single large batch of NGN2 neurons (∼7 billion), a viable screening window (Z’≥0.4) was maintained throughout the campaign, with two imaging rounds flagged for repeat testing (17 plates). Three methods were used to select hits, in order to maximize detection of diverse mechanisms. 1) Reversal of LDA disease score used in the HTS screen, 2) functional MPO score, in which individual features included in the LDA disease score were converted to z-scores and summed, and 3) recovery to wild type, in which compounds for which every feature included in the LDA disease score was within 3 standard deviations of the mean of the *TSC2* WT controls. Method 1 identified compounds that produced maximal reversal of disease score, while methods 2 and 3 were employed to select compounds that display a balanced form of phenotypic recovery, in which compound ranking is not driven by a smaller set of functional parameters.

### Hit characterization using pS6 high-content imaging

For additional characterization of hits from the phenotypic screen, immunocytochemistry (ICC) followed by high-content imaging was used to assess pS6 expression. ICC was carried out as described above. pS6 expression for each cell in a field-of-view was estimated using a custom automated image analysis package written in MATLAB. Briefly, image analysis consisted of using the human nuclear-specific immunoreactivity channel to identify pixels masks corresponding to individual cellular nuclei. Nuclear masks were expanded to create cytoplasmic masks that correspond to the neuronal cell body using the MAP2 channel. pS6 expression was quantified for each cell as the mean pixel intensity in the pS6 channel within the cell body mask for each cell. For screening analysis, single-cell expression values were aggregated over all cells in an experimental well. Assay wells on a 96-well plate were normalized by subtracting the mean of positive controls (TSC2-WT-3 neurons, 0.2% DMSO) and then dividing by the mean of the negative controls (TSC2-KO-112 cells, 0.2% DMSO) so that a normalized value of 0 represents the average pS6 expression in healthy control neurons and a normalized value of 1 represents the expression in the *TSC2* KO model.

### Hit characterization using acute vs. chronic treatment regimens

TSC2-KO-112 NGN2 neurons were treated with a set of 210 confirmed hits from the phenotypic screen. Compounds were dosed at 5 different concentrations (3x serial dilution: 10, 3.333, 1.111, 0.370, and 0.123 μM) using either the chronic treatment regimen used during the primary screening (**Figure 6A**, DIV10+24+31) or an acute dosing 30 to 90 minutes prior to optical physiology imaging. We classified a compound as active in a given treatment regimen if the average LDA score of wells treated with 3.333 μM of that compound was above 0.3. This LDA threshold was based on the behavior of *TSC2* KO and *TSC2* WT neurons treated with 0.1% DMSO. Using an activity threshold of 0.3, 98.8% of TSC2-KO + 0.1% DMSO wells (n=1568) were classified as inactive and 100% of TSC2-WT + 0.1% DMSO wells (n=1695) in this dataset were classified as active. We chose to evaluate activity based on LDA score after administration of 3.333 μM of drug because it is similar to the concentration used during the primary screening (5 μM). Principal component analysis (**Figure 7E**) in R was performed on functional data from DMSO-treated control neurons [*TSC2* WT (n=999 wells) and *TSC2* KO (n=872 wells)], *TSC2* KO neurons chronically treated with 0.3μM Torin-1 (n=79 wells), 30 μM XE-991 (n=64 wells), or 3.333 μM of the hit compounds that were active chronically or both chronically and acutely (n=2 replicate wells/group, defined in **Figure 7C**) using the *prcomp* function after normalization with scale. Experimental wells with missing data were removed. Similar analyses were carried out when *TSC2* KO neurons were treated with the different Kv channel modulators at different concentrations.

## ACKNOWLEDGMENTS

This work was funded by UCB Pharma and Quiver Bioscience NIH grants R44MH112474 and R44MH112273.

## AUTHOR CONTRIBUTIONS

G.T.D, C.W., L.A.W., S.J.R., A.E., and O.M. designed the study and provided overall scientific oversight. L.A.W. and V.J. designed and executed CRISPR/Cas9, iPS cell culture, neuronal production and immunocytochemistry experiments. V.J. and S.S. executed neuronal culture for disease modeling assays and primary phenotypic screen. J.G. and J.J. acquired neuronal functional data using Quiver’s all-optical physiology platform. J.J.F. designed and carried out preliminary synaptic assays. H.Z. designed and optimized 96-well functional measurements for primary screen. S.J.R., C.L., L.A.W., A.E., O.M, and G.T.D. analyzed and interpreted functional data. P.G. and C.L. carried out RNA-Seq analyses. M.G., Y.S., A.E. and O.M. performed selection of focused compound libraries for primary phenotypic screen. C.P., M.G., V.A., L.S., S.D. and C.W. provided scientific oversight related to *in vitro* screening activities and selection of small molecule compounds and confirmed hits. O.D. recruited and obtained TSC patient and healthy control blood samples for TSC iPS cellular reagent generation. L.A.W., A.E., G.T.D., and C.W. managed study timeline and budget. L.A.W., S.J.R., C.L., A.E., O.M., and G.T.D. prepared the manuscript. All authors contributed to manuscript revisions and approved the submitted version.

---

**Supplemental Figure 1.**
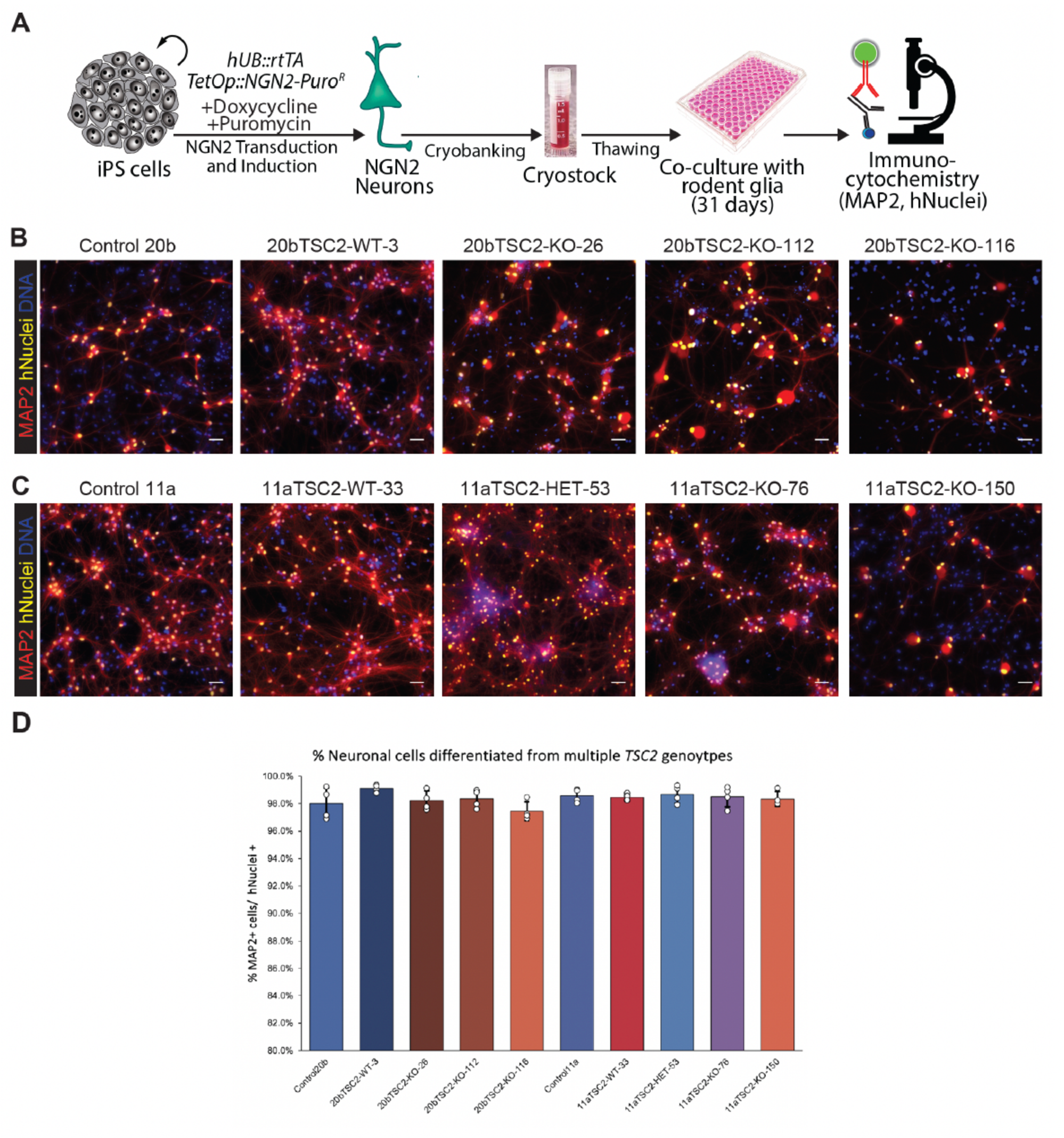
Production of NGN2 neurons from multiple CRISPR/Cas9-isogenic *TSC2* iPS cell lines. **(A)** Generation of cortical excitatory neurons via transcriptional programming with pro-neuronal factor NEUROG2 (NGN2). Neuronal cultures were characterized using immunocytochemistry (ICC) to determine fraction of human neurons in the preparations after co-culture with rodent primary glia for 31 days. **(B-C)** Fluorescence images of NGN2 neurons differentiated from multiple CRISPR/Cas9-isogenic *TSC2* iPS cell lines in two genetic backgrounds: 20b and 11a. Cultures were immunostained for the pan-neuronal marker MAP2 (Microtubule Associated Protein 2) in red, a human-specific nuclear antigen (hNuclei) in yellow, and DNA in blue. **(D)** Quantification of the ICC images showed that > 97% of human differentiated cells (hNuclei+) in the cultures expressed MAP2, indicating successful production of neurons independent of *TSC2* genotype.

**Supplemental Figure 2.**
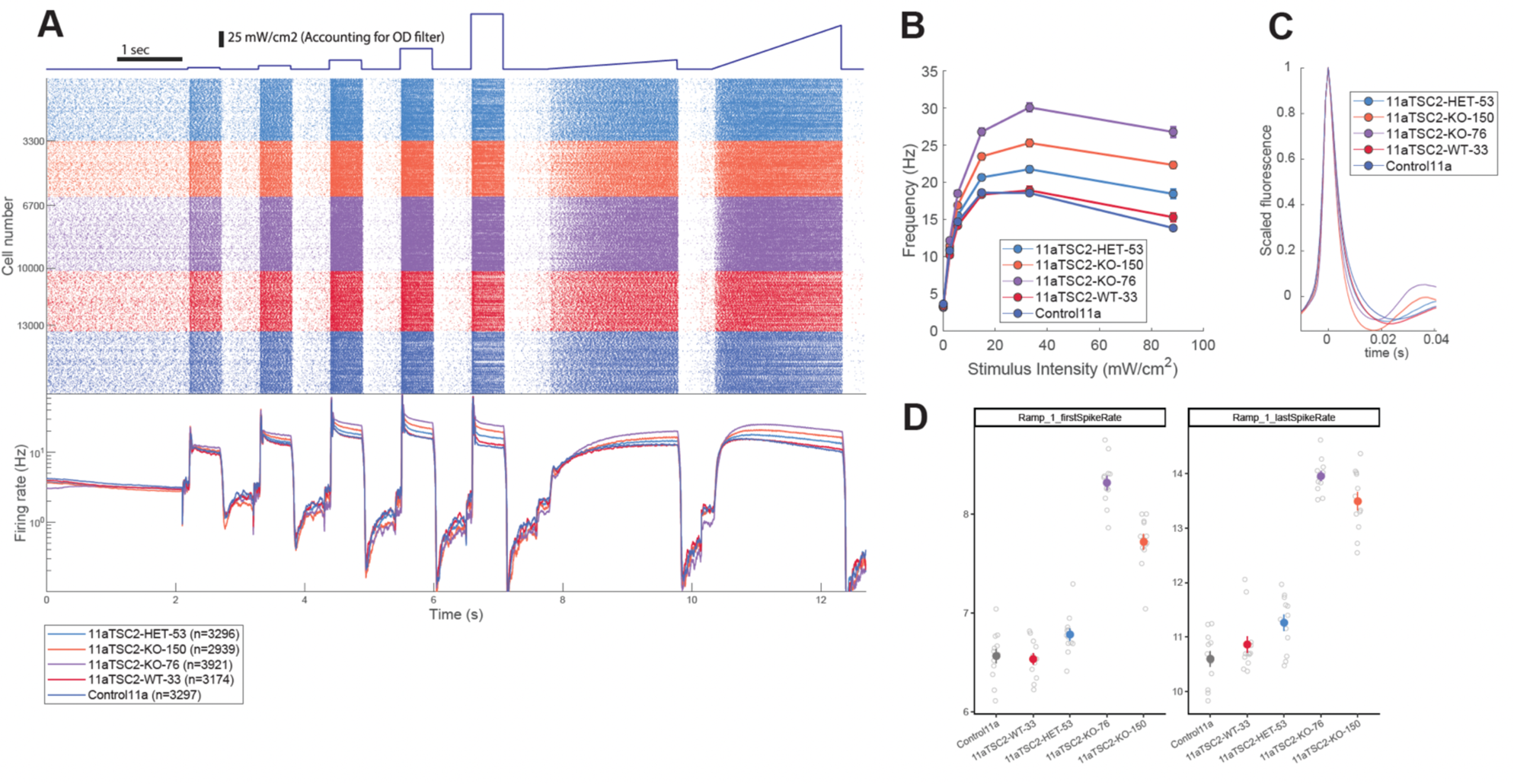
Functional phenotype of TSC2 loss is replicated in NGN2 neurons differentiated from the additional iPS cell genetic background 11a. **(A)** Functional characterization of NGN2 neurons obtained from TSC2 CRISPR/Cas9-isogenic iPS cell lines from 11a genetic background, including the parental genotype (Control 11a) and four clonal cell lines with disruption of the TSC2 gene after CRISPR/Cas9-mediated targeting: 11a TSC2-WT-33 clonal cell line is genetically identical to the parental line; 11a TSC2-HET-53 clonal cell line has a heterozygous *loss-of-function* mutation in *TSC2*, and 11a TSC2-KO-76 and 11a TSC2-KO-150 represent two cell lines with bi-allelic (homozygous) lessions in *TSC2*. Raster plot representing measurements of spike trains from >16,000 neurons (n = individuals neurons per genotype) and showing differential response to blue light stimulation according to *TSC2* genotype. **(B-D)** Neurons differentiated from 11a iPS cell genetic background showed a similar functional response to loss of TSC2 as seen in the 20b iPS cell background (**Figure 2**). Specifically, *TSC2* KO neurons fired more under high-intensity stimulation (**B** and **D**) and exhibited narrower action potentials (**C**). The heterozygous *TSC2* mutant genotype (11a TSC2-HET-53) exhibited a milder phenotype than *TSC2* KO neurons, with functional properties positioned in between *TSC2* WT and *TSC2* KO neurons for metrics related to spike timing.

**Supplemental Figure 3.**
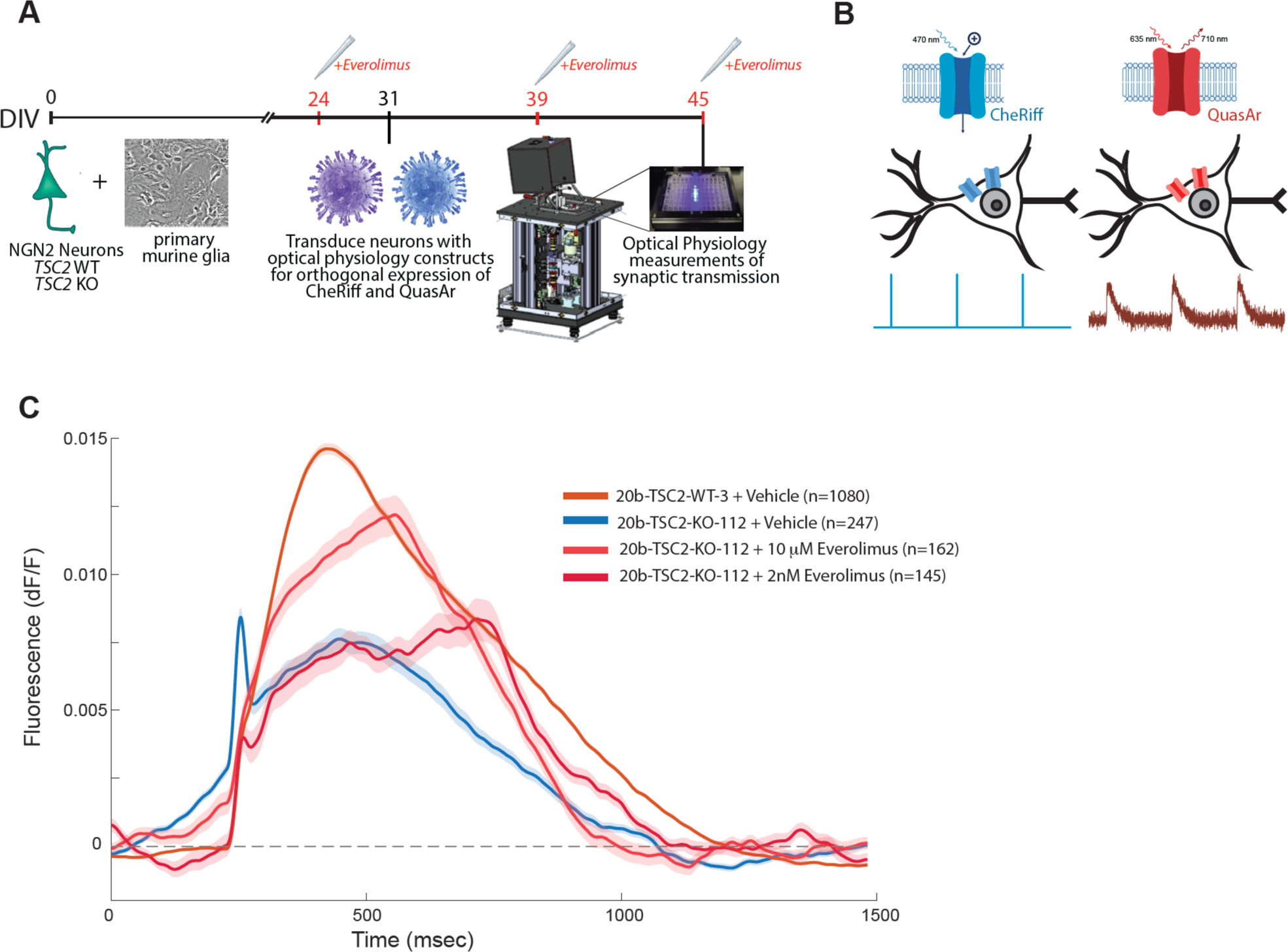
Synaptic phenotype and pharmacological rescue in *TSC2* KO. **(A)** Assay timeline: Excitatory NGN2 neurons from 20b-TSC2-WT-3 (*TSC2* WT) and 20b-TSC2-KO-112 (*TSC2* KO) iPS cell lines were were plated (DIV0, day *in vitro* 0) and co-cultured with rodent primary astrocytes and synaptic phenotype and pharmacological rescue were assessed on DIV45 using optical physiology measurements. Neurons were treated with either 2 nM or 10 µM Everolimus using a chronic and multi-intervention treatment regime (treatment at DIV24, DIV39, and acutely at DIV45 ∼1hr prior (and during) functional imaging. **(B)** Schematic of synaptic imaging protocol. To assess synaptic function, a Cre-based recombination approach was used to allow for orthogonal expression of the two optical physiology components, resulting in two distinct neuronal populations: those expressing the channelrhodopsin CheRiff, and those expressing the voltage reporter QuasAr. Three pulses of blue light were delivered to the neurons (as depicted), resulting in opening of the channelrhodopsin, depolarization of the membrane of CheRiff-expressing neurons, and subsequent action potential firing and synaptic transmitter release. The presence of QuasAr in the remaining neurons allowed for detection of changes in membrane voltage resulting from postsynaptic potentials due to neurotransmitter binding to glutamate receptors (GABAzine was included in imaging buffer to block inhibitory signaling). **(C)** Excitatory postsynaptic potential waveforms (expressed as changes to the fluorescence of the voltage reporter QuasAr) under vehicle (0.1% DMSO) or Everolimus conditions for *TSC2* KO vs. *TSC2* WT neurons. *TSC2* KO neurons showed a dramatically reduced postsynaptic potential that is partially rescued with increasing concentrations of the mTOR modulator Everolimus. 24 culture wells were measured per condition and n = the number of individual neurons in which a postsynaptic potential was detected. Note that many fewer *TSC2* KO neurons showed postsynaptic potentials as compared to *TSC2* WT despite an equal number of samples across conditions.

**Supplementary Table 1:**
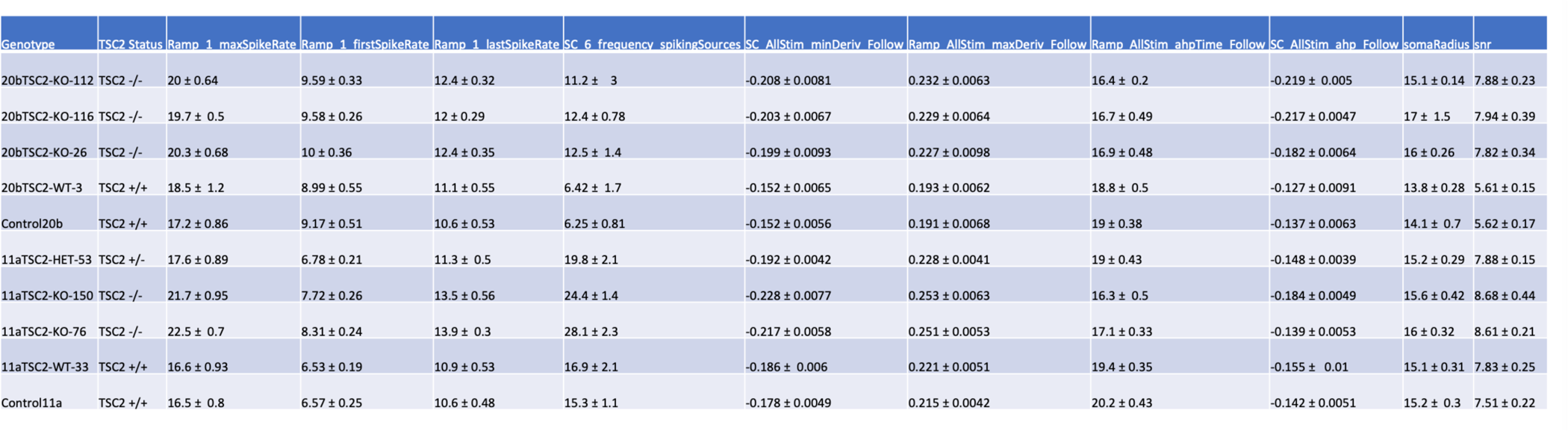
Functional phenotypic feature values for CRISPR/Cas9 *TSC2* isogenic cell lines.

**Supplementary Table 2:**
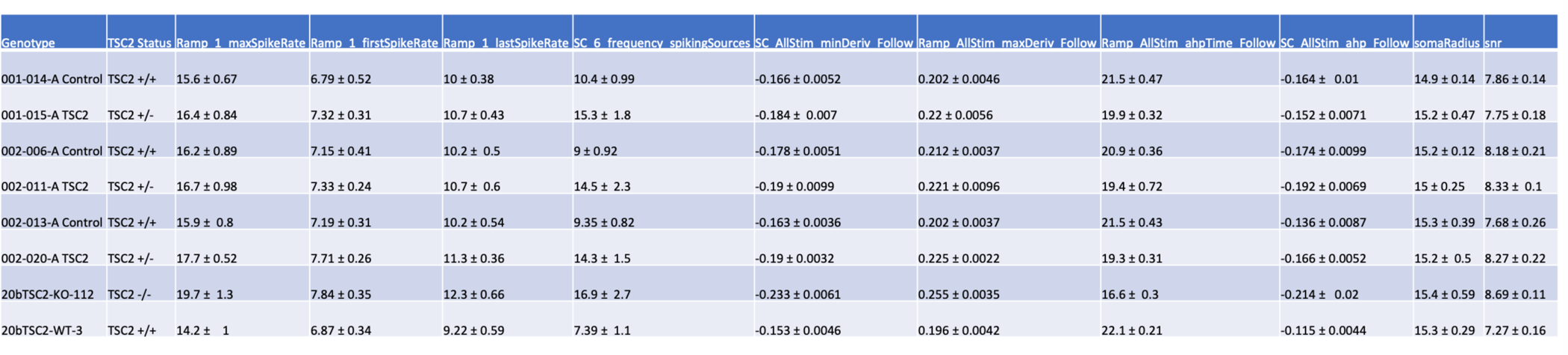
Functional phenotypic feature values for TSC Patient and Control cell lines.

## REFERENCES

1. Dowden, H. & Munro, J. Trends in clinical success rates and therapeutic focus. Nat. Rev. Drug Discov. 18, 495–496 (2019).

2. Lee, C. E., Singleton, K. S., Wallin, M. & Faundez, V. Rare Genetic Diseases: Nature’s Experiments on Human Development. iScience 23, 101123 (2020).

3. Condò, I. Rare Monogenic Diseases: Molecular Pathophysiology and Novel Therapies. Int. J. Mol. Sci. 23, 6525 (2022).

4. Simkin, D. et al. ‘Channeling’ therapeutic discovery for epileptic encephalopathy through iPSC technologies. Trends Pharmacol. Sci. 43, 392–405 (2022).

5. Ellis, C. A., Petrovski, S. & Berkovic, S. F. Epilepsy genetics: clinical impacts and biological insights. Lancet Neurol. 19, 93–100 (2020).

6. Lou, S. & Cui, S. Drug Treatment of Epilepsy: From Serendipitous Discovery to Evolutionary Mechanisms. Curr. Med. Chem. 29, 3366–3391 (2022).

7. Cook, A. M. & Bensalem-Owen, M. K. Mechanisms of action of antiepileptic drugs. Therapy 8, 307–313 (2011).

8. Strzelczyk, A. & Schubert-Bast, S. Psychobehavioural and Cognitive Adverse Events of Anti-Seizure Medications for the Treatment of Developmental and Epileptic Encephalopathies. CNS Drugs 36, 1079–1111 (2022).

9. Hyman, M. H. & Whittemore, V. H. National Institutes of Health consensus conference: tuberous sclerosis complex. Arch. Neurol. 57, 662–665 (2000).

10. Uysal, S. P. & Şahin, M. Tuberous sclerosis: a review of the past, present, and future. Turk. J. Med. Sci. 50, 1665–1676 (2020).

11. Curatolo, P., Specchio, N. & Aronica, E. Advances in the genetics and neuropathology of tuberous sclerosis complex: edging closer to targeted therapy. Lancet Neurol. 21, 843–856 (2022).

12. Overwater, I. E. et al. A randomized controlled trial with everolimus for IQ and autism in tuberous sclerosis complex. Neurology 93, e200–e209 (2019).

13. Nabbout, R., Kuchenbuch, M., Chiron, C. & Curatolo, P. Pharmacotherapy for Seizures in Tuberous Sclerosis Complex. CNS Drugs 35, 965–983 (2021).

14. de Wit, D. et al. Everolimus pharmacokinetics and its exposure-toxicity relationship in patients with thyroid cancer. Cancer Chemother. Pharmacol. 78, 63–71 (2016).

15. European Medicine Agency (EMA). (2010) Afinitor® (everolimus): European Public Assessment Report (EPAR). http://www.ema.europa.eu/.

16. Loewa, A., Feng, J. J. & Hedtrich, S. Human disease models in drug development. Nat. Rev. Bioeng. (2023) doi:10.1038/s44222-023-00063-3.

17. Costa, V. et al. MTORC1 Inhibition Corrects Neurodevelopmental and Synaptic Alterations in a Human Stem Cell Model of Tuberous Sclerosis. Cell Rep. 15, 86–95 (2016).

18. Li, Y. et al. Abnormal Neural Progenitor Cells Differentiated from Induced Pluripotent Stem Cells Partially Mimicked Development of {TSC2} Neurological Abnormalities. Stem Cell Rep. 8, 883–893 (2017).

19. Zucco, A. J. et al. Neural progenitors derived from Tuberous Sclerosis Complex patients exhibit attenuated {PI3K}/{AKT} signaling and delayed neuronal differentiation. Mol. Cell. Neurosci. 92, 149–163 (2018).

20. Sundberg, M. et al. Purkinje cells derived from TSC patients display hypoexcitability and synaptic deficits associated with reduced FMRP levels and reversed by rapamycin. Mol. Psychiatry 23, 2167–2183 (2018).

21. Nadadhur, A. G. et al. Neuron-Glia Interactions Increase Neuronal Phenotypes in Tuberous Sclerosis Complex Patient {iPSC}-Derived Models. Stem Cell Rep. 12, 42–56 (2019).

22. Winden, K. D. et al. Biallelic mutations in TSC2 lead to abnormalities associated with cortical tubers in human ipsc-derived neurons. J. Neurosci. 39, (2019).

23. Afshar Saber, W. & Sahin, M. Recent advances in human stem cell-based modeling of Tuberous Sclerosis Complex. Mol. Autism 11, 16 (2020).

24. Williams, L. A. et al. Scalable Measurements of Intrinsic Excitability in Human iPS Cell-Derived Excitatory Neurons Using All-Optical Electrophysiology. Neurochem. Res. 44, 714–725 (2019).

25. Werley, C. A., Chien, M.-P. & Cohen, A. E. An ultrawidefield microscope for high-speed fluorescence imaging and targeted optogenetic stimulation. Biomed. Opt. Express 8, 5794 (2017).

26. Boulting, G., Kiskinis, E. & Croft, G. A functionally characterized test set of human induced pluripotent stem cells. Nat. … 29, 279–286 (2011).

27. Zhang, Y. et al. Rapid Single-Step Induction of Functional Neurons from Human Pluripotent Stem Cells. Neuron 78, 785–798 (2013).

28. Bassetti, D., Luhmann, H. J. & Kirischuk, S. Effects of Mutations in TSC Genes on Neurodevelopment and Synaptic Transmission. Int. J. Mol. Sci. 22, (2021).

29. Hochbaum, D. R. et al. All-optical electrophysiology in mammalian neurons using engineered microbial rhodopsins. Nat. Methods 11, 825–833 (2014).

30. Werley, C. A. et al. All-optical electrophysiology for disease modeling and pharmacological characterization of neurons. Curr. Protoc. Pharmacol. 2017, 11.20.1–11.20.24 (2017).

31. Fisher, R. A. The Use of Multiple Measurements in Taxonomic Problems. Ann. Eugen. 7, 179–188 (1936).

32. Discriminant Analysis and Statistical Pattern Recognition | Wiley. Wiley.com https://www.wiley.com/en-us/Discriminant+Analysis+and+Statistical+Pattern+Recognition-p-9780471691150.

33. Niu, W., Siciliano, B. & Wen, Z. Modeling tuberous sclerosis complex with human induced pluripotent stem cells. World J. Pediatr. WJP (2022) doi:10.1007/s12519-022-00576-8.

34. Sandoe, J. & Eggan, K. Opportunities and challenges of pluripotent stem cell neurodegenerative disease models. Nat. Neurosci. 16, 780–9 (2013).

35. Bateup, H. S. et al. Excitatory/Inhibitory Synaptic Imbalance Leads to Hippocampal Hyperexcitability in Mouse Models of Tuberous Sclerosis. Neuron 78, 510–522 (2013).

36. Abs, E. et al. TORC1-dependent epilepsy caused by acute biallelic *Tsc1* deletion in adult mice: Epilepsy upon *Tsc1* Deletion. Ann. Neurol. 74, 569–579 (2013).

37. Vincent, F. et al. Phenotypic drug discovery: recent successes, lessons learned and new directions. Nat. Rev. Drug Discov. 21, 899–914 (2022).

38. Di Giorgio, F. P., Boulting, G. L., Bobrowicz, S. & Eggan, K. C. Human Embryonic Stem Cell-Derived Motor Neurons Are Sensitive to the Toxic Effect of Glial Cells Carrying an ALS-Causing Mutation. Cell Stem Cell 3, 637–648 (2008).

39. Williams, L. A. et al. Developing antisense oligonucleotides for a TECPR2 mutation-induced, ultra-rare neurological disorder using patient-derived cellular models. Mol. Ther. Nucleic Acids 29, 189–203 (2022).

40. Borja, G. B. et al. Probing Synaptic Signaling with Optogenetic Stimulation and Genetically Encoded Calcium Reporters. in 109–134 (2021). doi:10.1007/978-1-0716-0830-2_8.

41. Mukamel, E. A., Nimmerjahn, A. & Schnitzer, M. J. Automated Analysis of Cellular Signals from Large-Scale Calcium Imaging Data. Neuron 63, 747–760 (2009).

42. Benjamini, Y. & Hochberg, Y. Controlling the False Discovery Rate: A Practical and Powerful Approach to Multiple Testing. J. R. Stat. Soc. Ser. B Stat. Methodol. 57, 289–300 (1995).

43. Patro, R., Duggal, G., Love, M. I., Irizarry, R. A. & Kingsford, C. Salmon provides fast and bias-aware quantification of transcript expression. Nat. Methods 14, 417–419 (2017).

44. Soneson, C., Love, M. I. & Robinson, M. D. Differential analyses for RNA-seq: transcript-level estimates improve gene-level inferences. F1000Research 4, 1521 (2015).

45. Robinson, M. D. & Oshlack, A. A scaling normalization method for differential expression analysis of RNA-seq data. Genome Biol. 11, R25 (2010).

46. Johnson, W. E., Li, C. & Rabinovic, A. Adjusting batch effects in microarray expression data using empirical Bayes methods. Biostat. Oxf. Engl. 8, 118–127 (2007).

47. Law, C. W., Chen, Y., Shi, W. & Smyth, G. K. voom: Precision weights unlock linear model analysis tools for RNA-seq read counts. Genome Biol. 15, R29 (2014).

48. Liu, R. et al. Why weight? Modelling sample and observational level variability improves power in RNA-seq analyses. Nucleic Acids Res. 43, e97 (2015).

49. Hu, Y., Lounkine, E. & Bajorath, J. Improving the search performance of extended connectivity fingerprints through activity-oriented feature filtering and application of a bit-density-dependent similarity function. ChemMedChem (2009) doi:10.1002/cmdc.200800408.

50. Rogers, D. & Hahn, M. Extended-connectivity fingerprints. J. Chem. Inf. Model. (2010) doi:10.1021/ci100050t.

51. Wager, T. T., Hou, X., Verhoest, P. R. & Villalobos, A. Moving beyond rules: The development of a central nervous system multiparameter optimization (CNS MPO) approach to enable alignment of druglike properties. ACS Chem. Neurosci. (2010) doi:10.1021/cn100008c.

52. Baell, J. B. & Holloway, G. A. New substructure filters for removal of pan assay interference compounds (PAINS) from screening libraries and for their exclusion in bioassays. J. Med. Chem. 53, 2719–2740 (2010).

